# Effects of noncanonical genomic imprinting in monoaminergic pathways on the regulation of social behaviors

**DOI:** 10.1101/2024.02.28.582619

**Authors:** Erin M. O’Leary, Samuel J. Rahman, Andrei L. Tamas, Tony Huang, Moudar Dweydari, Rachel L. Eggleston, Daryl D. Meling, Paul J. Bonthuis

## Abstract

Genomic imprinting in the brain is theorized to provide parental control over offspring social behaviors. Noncanonical genomic imprinting is a form of epigenetic regulation in which one of a gene’s alleles, either that of maternal or paternal inheritance, exhibits a bias towards higher expression of one parental allele compared to the other. This bias can differ depending on tissue type, and the degree of the parental allele expression bias can even vary across anatomical domains within the same tissue. Dopa decarboxylase (*Ddc*) and tyrosine hydroxylase (*Th*) are both noncanonically imprinted genes that preferentially express their maternal alleles in the brain and *Ddc* also has a paternal allele expression bias in the periphery. These two genes encode catecholamine synthesis enzymes for the production of dopamine (DA), norepinephrine (NE), and epinephrine (E), and *Ddc* is also in the serotonin (5-HT) synthesis pathway. These four neurotransmitters are critical regulators of social behavior and disruptions to them are implicated in human mental illnesses. Here we investigated the functional effects of noncanonical imprinting of *Ddc* and *Th* on social behavior in mice. By using reciprocal heterozygous mutant mice, we tested the impacts of *Ddc* and/or *Th* maternally and paternally inherited alleles on aggression, social recognition, dominance, and social preference behaviors. We found that *Ddc* paternal-null alleles affect aggression and social recognition behavior, *Th* maternal-null alleles affect sociability preferences, and compound inheritance of *Th* and *Ddc* maternal-null alleles influence preferences for social novelty. These results are consistent with *Th* and *Ddc* maternal allele biased expression in central monoaminergic systems regulating sociability, and *Ddc* paternal allele biased expression in peripheral monoaminergic systems regulating aggression and social recognition.

## 1 | Introduction

The expression of imprinted genes is highly prevalent in the brain, and genomic imprinting theories and empirical evidence from humans and animal models indicate that imprinting effects in the brain have evolved for parental control over offspring social behaviors (Keverne et al., 1996; Isles et al., 2006; Barlow and Bartolomei, 2014; McNamara and Isles, 2014; Bonthuis et al., 2015; Andergassen et al., 2017; Lassi and Tucci, 2017; McNamara et al., 2018b; Higgs et al., 2022). Canonical genomic imprinting is an epigenetic process in mammals and certain flowering plants, by which either the maternally or paternally inherited allele of an imprinted gene is exclusively expressed, while the allele from the other parent is silenced in offspring (Bartolomei and Ferguson-Smith, 2011; Ho-Shing and Dulac, 2019). Human genetic disorders involving imprinted genes, including Angelman syndrome (AS) and Prader-Willi syndrome (PWS), result in atypical neurodevelopment that can affect behavioral traits relevant to social interaction (Horsler and Oliver, 2006; Isles et al., 2006; Cassidy and Driscoll, 2009; Ho-Shing and Dulac, 2019). A mouse model of AS with a maternally inherited heterozygous deletion of *Ube3a* shows impaired learning and increased ultrasonic vocalizations indicative of abnormal social communication (Jiang et al., 2010) and another mouse model with a mutation at the AS/PWS locus with overexpression of the paternal allele displayed impaired social interactions reflective of human autism (Nakatani et al., 2009). Mice with a paternal mutation of *Magel2*, which is in the PWS deletion region, also have phenotypes that mirror the human disorder, including altered feeding behavior, increased anxiety, reduced fertility, and disrupted circadian rhythms (Mercer and Wevrick, 2009; Bervini and Herzog, 2013). Mouse studies have also leveraged mutant models to examine the roles of particular imprinted genes in physiology and behavior. For example, paternal allele expression of *Peg3* has been found to regulate maternal behavior and offspring growth (Li et al., 1999; McNamara et al., 2018a) and paternal allele expression of *Grb10* regulates impulsive behavior and dominance (Garfield et al., 2011; Dent et al., 2020). Thus far, most evidence examining the role of imprinted genes in social behavior focuses on *canonically* imprinted genes where mutations cause either a loss of the expressed allele or biallelic expression of a gene normally only expressed from one allele (Li et al., 1999; Jiang et al., 2010; Creeth et al., 2018; Dent et al., 2018; McNamara et al., 2018a; Higgs et al., 2023).

Recent work has utilized genome-wide RNA sequencing techniques to measure allelic expression profiles across the whole genome to characterize the extent of genomic imprinting effects throughout the body (Laukoter et al., 2020; Varrault et al., 2020; Edwards et al., 2023). These studies demonstrate that imprinting effects can be highly tissue-specific, are enriched in the brain, and identify noncanonically imprinted allelic expression effects in addition to canonical imprinting effects. (Gregg et al., 2010; Bonthuis et al., 2015; Perez et al., 2015; Andergassen et al., 2017). In contrast to canonical genomic imprinting consisting of allelic silencing of one parental allele and monoallelic expression of either a maternally expressed gene (MEG) or paternally expressed gene (PEG), noncanonical imprinting effects represent a consistent tissue-level bias to express the maternal or paternal allele to a higher level than the other allele (Bonthuis et al., 2015; Huang et al., 2018). However, few studies have demonstrated the functional implications for noncanonical imprinting effects in the brain or other tissues.

RNAseq studies found a network of noncanonical imprinted monoamine neurotransmitter synthesis and signaling genes in the brain (Bonthuis et al., 2015). Monoaminergic systems releasing dopamine (DA), serotonin (5-HT), norepinephrine (NE), and epinephrine (E) have well-known neuromodulatory activities involved in a wide range of animal behaviors that include reward, anxiety, stress, and social behavior (Morilak et al., 2005; Belujon and Grace, 2015; Brummelte et al., 2017; Wise and Jordan, 2021). Two noncanonically imprinted monoamine neurotransmitter synthesis genes, tyrosine hydroxlyase (*Th*) and dopa decarboxylase (*Ddc*), were found to preferentially express the maternal allele in certain regions of the brain (Bonthuis et al., 2015, 2022). *Th* is required for catecholamine biosynthesis, which includes DA, NE, and E (Daubner et al., 2011). *Ddc* is required for catecholamine and serotonin 5-HT(Bertoldi, 2014) biosynthesis (Fig. 1a). In pyrosequencing allelic expression studies, both *Th* and *Ddc* were found to have a maternal allele expression bias in several distinct monoaminergic brain regions: namely the arcuate nucleus (ARN) of the hypothalamus (DA region), the dorsal raphe (DR) of the midbrain (5-HT region), and the locus coeruleus (LC) of the pons (NE region) (Bonthuis et al., 2015). Although the aforementioned study found no significant maternal allele expression bias for either gene in the dopaminergic ventral tegmental area (VTA) of the midbrain (Bonthuis et al., 2015), a more recent study found evidence for higher expression from the *Ddc* maternal allele compared to the paternal allele in midbrain sections using a *Ddc-hKO1* fluorescent reporter knock-in mouse line and fluorescence microscopy analysis (Sheng et al., 2022). Using reciprocal crosses of two different knock-in allelic reporters (*Ddc-P2A-6His-eGFP* and *Ddc-P2A-V5-mRuby2),* we found *Ddc* maternal monoallelic subpopulations were particularly present in the hypothalamus (Bonthuis et al., 2022), which has a vast number of molecularly distinct cell populations (Saper and Lowell, 2014; Chen et al., 2017; Chen and Hong, 2018; Kim et al., 2019). Together with nascent RNA *in situ* hybridization studies showing monoallelic expression of *Th* and *Ddc* in brain regions with maternal allele expression biases (Bonthuis et al., 2015), analysis of *Ddc* allelic reporter mice indicated that noncanonical imprinting effects at the tissue level represents a mixture of highly cell-type specific imprinting effects and biallelic expression at the cellular level (Bonthuis et al., 2022). *Ddc* is also known to be imprinted in peripheral tissues other than the brain; it is paternally expressed in embryonic heart tissue, and in the medulla of the adrenal gland (MenhenioE et al., 2008; Babak et al., 2015; Chan et al., 2019; Bonthuis et al., 2022). Chromaffin cells in the adrenal medulla secrete E and NE into the bloodstream as a part of the sympatho adrenomedullary system (Eiden and Jiang, 2018). Thus, imprinting effects for some genes critical for monoaminergic systems are highly plastic with tissue and cell-type specificity.

**Fig 1.**
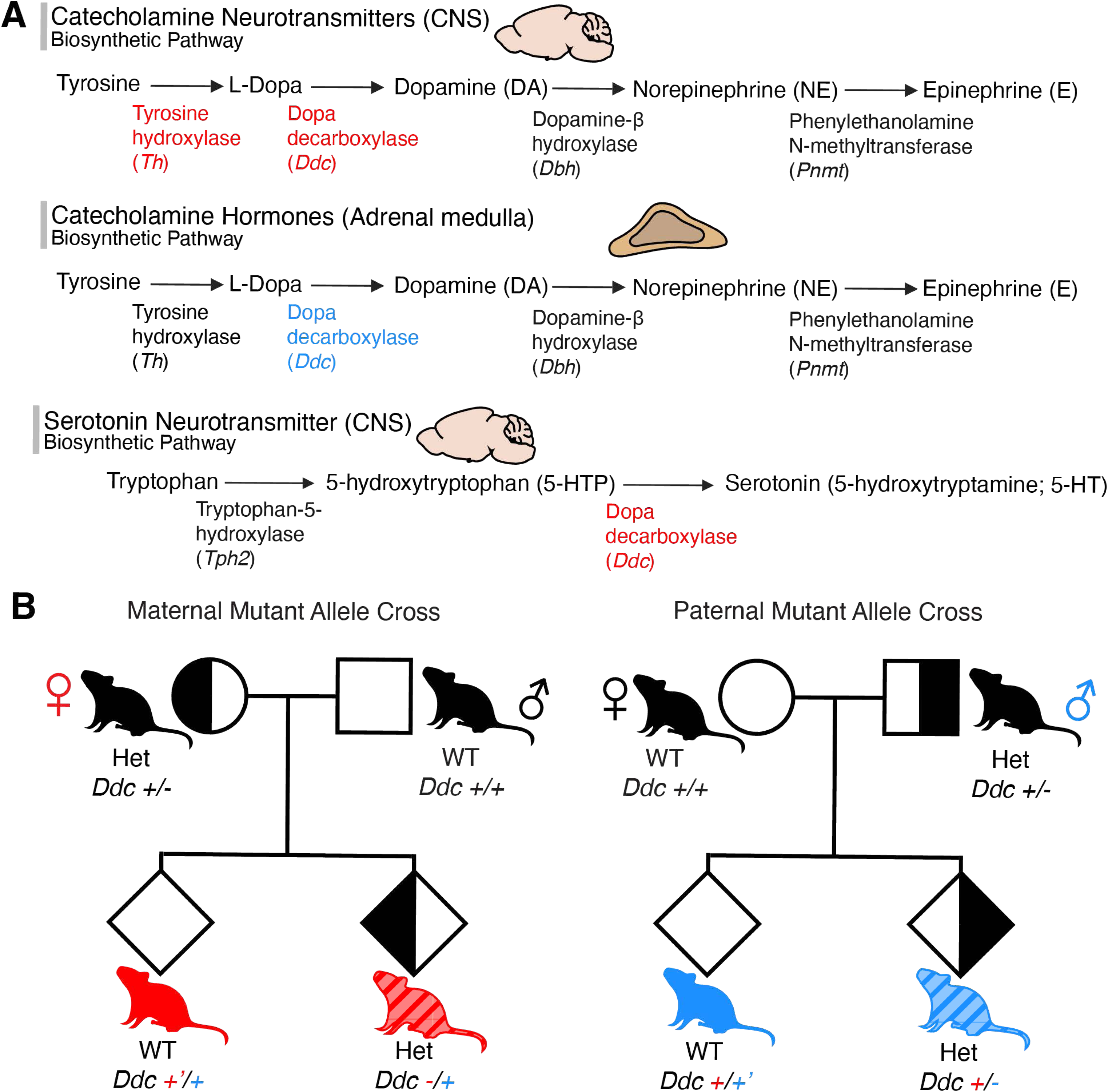
Dopa decarboxylase and Tyrosine hydroxylase are noncanonically imprinted genes in the catecholamine and serotonin biosynthesis pathways. **(a)** Tyrosine hydroxylase (*Th)* and dopa decarboxylase (*Ddc*) are catecholamine synthesis pathway genes and are noncanonically imprinted with maternally biased expression in the brain (red). *Ddc* has paternally biased expression in catecholamine producing cells of the adrenal medulla (blue). *Ddc* also synthesizes serotonin and has maternally biased expression in the brain (red). **(b)** *Ddc* maternal and paternal heterozygous mutant cross offspring from reciprocal WT and *Ddc* heterozygous parents. Offspring genotypes: solid red *Ddc* +’/+, maternal WT; striped red *Ddc* -/+, maternal heterozygote; solid blue *Ddc* +/+’, paternal WT; striped blue *Ddc* +/-, paternal heterozygote.

*Th* is well known for its role as the rate limiting step to catecholamine neurotransmitter synthesis (Flatmark, 2000; Daubner et al., 2011); and there is evidence for region-specific *Th* imprinting in the brain (Bonthuis et al., 2015). *Th* is located near several other well-known imprinted genes, including insulin-like growth factor 2 (*Igf2*) and *H19* within an imprinted gene cluster of mouse chromosome 7 (Lenartowski and Goc, 2011). Clinical studies in humans have demonstrated paternal uniparental isodisomy at the *TH01* locus, where both *TH* alleles are inherited from chromosomes of the same parent, results in abnormal gestation and nonviable pregnancy (Buza et al., 2019). Functional studies using reciprocal *Th* heterozygous mice that inherit a germline mutation from either the mother or the father found evidence that genomic imprinting in the catecholamine system affects anxiety-like and hedonic behaviors (Bonthuis et al., 2015). Furthermore, in a naturalistic foraging task, it was found that maternally inherited *Th* and *Ddc* alleles regulate male offspring foraging behavior, while paternally inherited alleles regulate female foraging behavior (Bonthuis et al., 2022). Additionally, at the endocrine level germline mutation of the *Ddc* maternal allele affects basal systemic DA, NE, and E concentrations in males, while the paternal allele affects DA concentrations in females (Bonthuis et al., 2022).

Here, we sought to test for the first time the role of noncanonical imprinting of *Th* and *Ddc* on social behaviors. We utilized animals with reciprocal heterozygous *Th* and/or *Ddc* functionally null deletion alleles to reveal social behavioral phenotypes affected by maternally inherited alleles compared to paternally inherited alleles. Behavioral differences between mice with a null maternal allele (-/+) to their wildtype littermates (+’/+) reveals traits that are regulated by the maternal allele. Conversely, traits affected by the paternal alleles are discovered by differences in mice with a null paternal allele (+/-) when compared to their wildtype (+/+’) littermates. In a first set of experiments, we look at the effects of maternally and paternally inherited heterozygous *Ddc* null alleles on offspring social behaviors. We tested offspring from reciprocal maternal and paternal *Ddc* null allele crosses (Fig. 1B) in aggression, social recognition, and social dominance tasks. In these experiments, we found that *Ddc^+/-^* paternal-null allele heterozygous mutations increase aggressive responses and impair social recognition in male offspring. In a second set of experiments, we use offspring from reciprocal compound heterozygote (Fig. 5A) crosses to determine if social preference behaviors are affected by catecholamine pathway noncanonical imprinting of either *Th* and *Ddc* alone, or synergistically (*Th^-/+^Ddc^-/+^*maternal null vs. *Th^+/-^Ddc^+/-^* paternal null offspring). We found that heterozygous mutations of *Ddc* from either cross (maternal-null and paternal-null alleles) reduce social preferences and that *Th^-/+^* maternal-null, but not paternal-null, heterozygous alleles reduce social preference behaviors in males. In addition, compound *Th^-/+^Ddc^-/+^* maternal-null heterozygous mice indicated that noncanonical imprinting in the same pathway synergically amplify maternal allele effects on social novelty preference. These findings implicate that noncanonically imprinted maternal and paternal allele expression of monoamine synthesis genes functionally regulate distinct social behavior phenotypes.

## 2 | Methods

### 2.1 | Housing and husbandry

Behavior experiments with *Ddc* heterozygotes were conducted in compliance with protocols approved by the University of Illinois at Urbana-Champaign’s (UIUC) Institutional Animal Care and Use Committee (IACUC). *Ddc* heterozygous mice (see below) were bred and housed on ventilated racks on a 12hr reversed light cycle. All mice were given water and food (Teklad 2918) *ad libitum*. Adult breeders (6 weeks to 1 year of age) were paired continuously, and pups were weaned at postnatal day 21 (P21) and cohoused in groups of two to five same-sex littermates or similar aged same-sex mice from the same cross. As needed, ear punches were taken before weaning for both genotyping biopsy samples and mouse identification purposes.

Social preference and social novelty experiments were conducted in compliance with protocols approved by the University of Utah Institutional Animal Care and Use Committee (IACUC). *ThDdc* compound heterozygous mice (see below) were bred and housed on static racks on a 12hr reversed light cycle. All mice were given water and food (Teklad 2920X soy protein-free) *ad libitum*. Breeding, weaning, and ear punches were the same as at UIUC.

### 2.2 | *Th* and *Ddc* heterozygote mutants

Germline heterozygous *Ddc* null allele mutant mice were made by crossing CMV-Cre (Jax, Stock No: 006054) X *Aadc^flox7^* lines and backcrossed to C57Bl/6J background strain for more than 10 generations as described previously (Bonthuis et al., 2022). The germline *Th* knockout mice (*ThΔ)* were maintained in the C57Bl/6J background (Jackson et al., 2012; Bonthuis et al., 2015). *ThDdc* compound heterozygous mice were made by crossing *Ddc* and *Th* heterozygous mice.

### 2.3 | Genotyping

Ear punches were taken at P7-P21. Samples were either genotyped by Transnetyx (Cordova, TN) using real-time PCR or were lysed in 75µL of 25mM NaOH + 0.2mM EDTA with a 1-hour incubation in a thermalcycler at 98°C. Lysates were then pH neutralized with an equal volume of 40mM Tris-HCl, pH 8.0. One µL of lysate was then added to make 20µL PCR reactions with GoTaq G2 Green Master Mix (Promega, Cat# M7823) and 1µM forward and reverse primers (Table 1). PCR reaction thermal cycling conditions were conducted using the following protocol: initial denaturation at 95°C for 3 minutes; then denaturation at 95°C for 30 seconds, annealing at 61°C for 30 seconds, and extension at 72°C for 1 minute for 28 cycles; and a final extension at 72°C for 5 minutes. Amplicons were run on a 2% agarose gel.

**Table 1.**
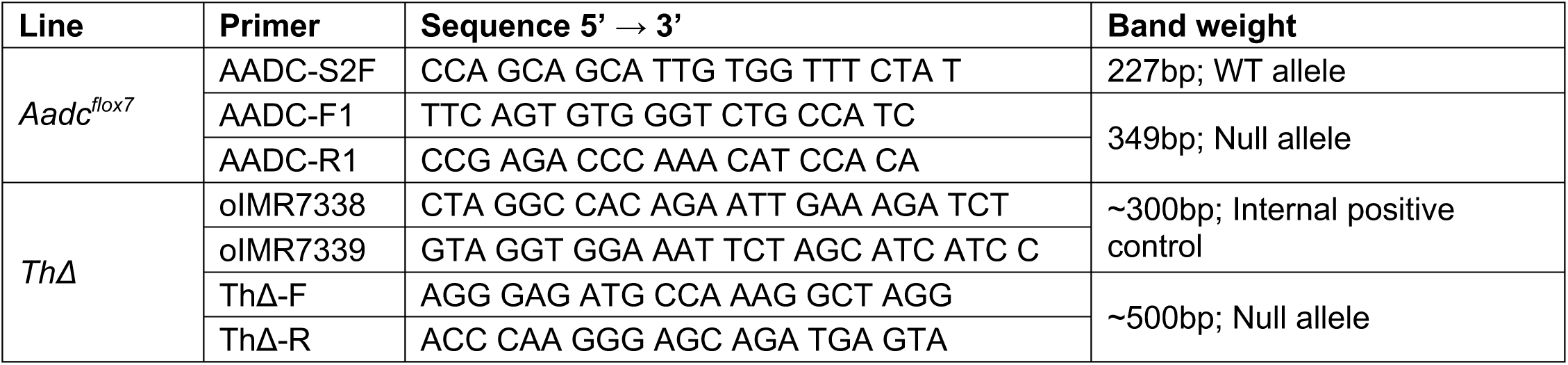
Genotyping primer sequences.

### 2.4 | Behavioral tests of reciprocal *Ddc* heterozygous mice

Behavioral tests of all reciprocal *Ddc* cross offspring genotype by sex groups (Fig. 1B) began between 8-10 weeks of age (P54-P75). Mice were tested once per week during the dark phase of the light:dark cycle. Bedding was left unchanged for 5–8 days prior to each test. Mice were transferred in their home cages and habituated to the testing room with the door closed and ambient room lights off for at least 30 minutes prior to any test. Testing occurred under red lighting with additional infrared lighting to properly illuminate the testing field. All tests were recorded with a digital camera (ELP, HD Infrared USB Webcam, 2.8 – 12 mm varifocal lens). One group of mice performed the social recognition test then aggression testing one week later. Mice tested for social dominance only performed that one test.

#### 2.4.1 | Social recognition test

Social recognition was assessed by use of a ten-trial social recognition task (Tejada and Rissman, 2012). Mice were housed individually for seven to fourteen days. On the day of testing mice were moved to the testing room and allowed to acclimate for at least one hour. Testing occurred in clear plexiglass boxes (30 cm x 18 cm x 14 cm) containing their home cage bedding; a round wire cup (7.6 cm diameter) was placed 4 cm from the edge of the box. Ten investigation trials, 1 minute each, were separated by 10-minute intertrial intervals. During the first nine trials, the same unfamiliar, same-sex stimulus mouse conspecific was placed in the home cage under a wire cylinder. On the tenth trial, a novel same-sex stimulus mouse was used. The total amount of time spent investigating the cylinder and/or the stimulus animal was recorded. Investigation was defined as nose contact with the stimulus mouse or wire cylinder at a distance of 2 cm or less. The avoidance zone (10 cm x 18 cm) was defined as the side of the cage opposite of the wire cup containing the conspecific. Ovariectomized C57Bl/6J females were used as stimuli for female test subjects and age matched C57Bl/6J males were used as stimuli for male test subjects. Stimulus animals were acclimated to the wire cylinder for three days prior to testing in three 1-minute sessions each day. All tests were tracked using DeepLabCut and behavioral measures were scored with SimBA (description of methods below). To assess habituation to repeated investigation of the same social stimulus, we analyzed the difference in investigation time for trials 2-9 compared to the first trial baseline and for the dishabituation trial we compared the difference between trial 9 and 10.

#### 2.4.2 | Resident-intruder test

Male aggression was measured with the resident-intruder test (Koolhaas et al., 2013). Male residents were housed with an intact female companion for at least one week prior to testing to increase aggressive behaviors (Albert et al., 1988). Bedding was not changed in the week prior to testing. On the day of testing the female companion was removed one hour before the start of testing. Males were moved to the testing room and allowed to acclimate for at least one hour. Intruders were from the A/J strain, chosen for their passive phenotype (Guillot and Chapouthier, 1996); in the few cases where intruders initiated aggression or retaliated after an aggressive attack, trials were excluded (*Ddc^+’/+^*, n = 3; *Ddc^-/+^* n = 1). An unfamiliar male intruder was introduced to the resident’s cage and interactions were recorded for 10 minutes. At the end of the test, the A/J intruder was removed, and the female companion was replaced. Testing continued for two more days, with a different A/J intruder male presented to the resident each day. Behavior was scored from video (Mouse Ethogram) and included latency to attack (boxing, aggressive bite, attack, fighting, chase) (s), number (n) and duration of attacks (s), social exploration (anogenital sniffing, social grooming, social exploration) (s), and nonsocial behavior (non-social exploration, resting, inactivity) (s). All tests were recorded and scored by an investigator with ANY-maze TakeNote (Stoelting; Wood Dale, Illinois) application.

#### 2.4.3 | Dominance tube test

Social dominance was measured with the tube test (Fan et al., 2019). Sex matched *Ddc^-/+^*, *Ddc^+/-^*, and WT siblings from each cross (*Ddc^+’/+^* maternal cross, and *Ddc^+/+’^* paternal cross) were housed together starting between P22-P35 to create a cage of four mice, one mouse per genotype (Tables 2, 3). Testing began when mice reached age P60. Two days of training were completed prior to beginning the test. Training consisted of teaching the mouse to walk through a clear polycarbonate tube 30 cm in length, 2 cm in diameter. Mice were followed by a plastic stick and were gently touched on their tails if they retreated or stopped moving. This was repeated ten times, alternating which end of the tube the mouse entered. On the day of the test, mice were moved to the testing room and allowed to acclimate for at least one hour. A round-robin style tournament was used to determine cage hierarchy. Each mouse faced off against all its cage-mates once per day, for a total of six face offs (three per mouse) each day. Pairs of cage-mates were presented to each other at either end of the clear polycarbonate tube. Their tails were released simultaneously, and the mice met in the middle of the tube; the first to retreat with all four paws out of the tube was designated the “loser”, and the other the “winner.” If no winner after 2 minutes, face-off was considered a draw. Both mice were returned to their home cage, the tube was cleaned with a 70% ethanol solution, and the next pair was faced off. At the conclusion of each day of testing mice were ranked by the number of wins against the other three cage-mates. Testing order was counter-balanced by genotype across days of testing. Once the mice maintained the same ranking for four consecutive days, rank was considered stable, and no further testing occurred. Final rank and average wins per day were measured for each mouse.

**Table 2.**
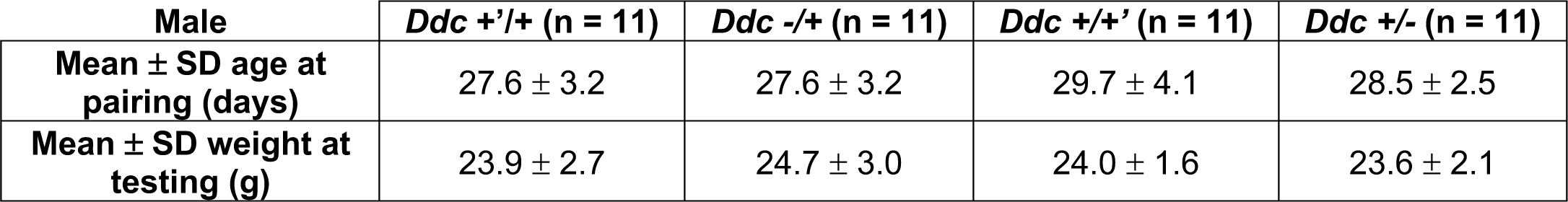
Male quartets average age at pairing and weight at testing by genotype.

**Table 3.**
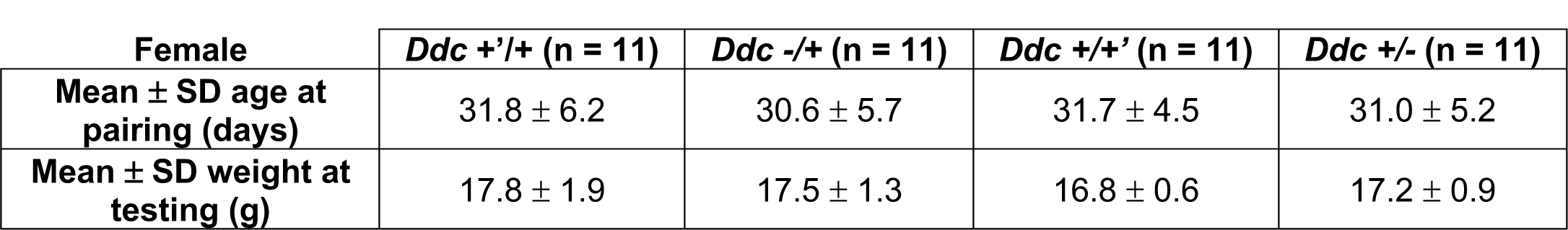
Female quartets average age at pairing and weight at testing by genotype.

### 2.5 | Compound *ThDdc* sociability and social novelty

Sociability and social novelty tests were performed with offspring from reciprocal *ThDdc* compound heterozygous crosses (Fig. 5a). Mouse movements were tracked with Ethovision XT 14 software (Noldus; Wageningen, Netherlands) under infrared illumination to collect behavioral data. Standard three-chambered boxes (Stoelting, Cat# 60450) with center, left, and right chambers separated by doors, were used to test social preference and social novelty behaviors. In the week prior to testing, social stimulus conspecific adult C57Bl/6 males were habituated to being confined to the round wire cups used in the three-chambered box for 30 minutes for 3-5 days. Tests were recorded in the dark with infrared illumination, and two mice were tested simultaneously in separate mazes. In the *sociability* task, one chamber (left or right) was occupied with a conspecific confined to a wire cup, while the opposite chamber (right or left) contained an empty wire cup. To begin the sociability task, the test mouse was habituated to the center chamber for five minutes with the doors closed to the left and right chambers. The experimenter then started Ethovision recording and removed the doors between the chambers to allow the test mouse to investigate for 10 minutes. Immediately following the 10-minute sociability test, the test mouse was returned to the center chamber with the doors closed to begin the *social novelty* task. For the social novelty task, a “novel” stranger conspecific was placed into the previously unoccupied wire cup while the now “familiar” conspecific used in the preceding social preference task remained in its’ wire cup. The experimenter then removed the doors while Ethovision recorded the test mouse investigating the chambers and wire cups with either the “familiar” or “novel” conspecific for an additional 10-minute trial. Conspecific stimulus mice were not reused in consecutive tests and use of the left or right chamber for the first/familiar and empty/novel conspecifics was counterbalanced by sex and genotype. Ethovision *Analysis Profile* exported the following measures for the *sociability* task: total distance moved (cm); maximum velocity (cm/s); cumulative duration (s), frequency of entries (n), mean duration per entry (s), and latency to enter (s) the conspecific chamber; cumulative duration (s), frequency of entries (n), mean duration per entry (s), and latency to enter (s) the empty chamber; cumulative duration (s), frequency of entries (n), and mean duration per entry (s) in the center chamber; cumulative duration (s), frequency (n), mean duration (s), and latency(s) for nose to investigate conspecific cup (s); cumulative duration (s), frequency (n), mean duration (s), and latency (s) for nose to investigate empty cup; and fecal boli (n). The same measures were collected for the social novelty task, except the chambers and cups contained either a “familiar” or “stranger” conspecific.

### 2.6 | Manual behavioral scoring

Social dominance and resident-intruder were all recorded and scored manually by a trained observer blind to genotype. Resident-intruder behaviors were scored using the TakeNote feature of ANY-maze (Stoelting; Wood Dale, Illinois).

### 2.7 | Automated behavioral scoring

A combination of a marker-less pose-estimation toolkit, DeepLabCut (DLC) (Mathis et al., 2018), and a region-of-interest and supervised behavior classifier pipeline, Simple Behavior Analysis (SimBA) (Nilsson et al., 2020), were used track mice and score behaviors in the social recognition test. DLC used a deep learning neural network trained on 420 manually annotated frames taken from 21 experimental videos. For this, an experimental observer annotated select frames for 8 body parts: the mouse’s snout, left ear, right ear, center midsection, left midsection, right midsection, tail base, and tail tip. The manually annotated frames were used by DLC pose-estimation algorithms to estimate the location of these body parts throughout all frames of the 1,360 1 min. social recognition videos. After training the DLC neural network for 250,000 iterations, the machine-labeled pose-estimations fell within an acceptable range of accuracy of 5 pixels compared to manually annotated frames: an effective accuracy of 3.45mm of physical space. DLC pose-estimation data was imported to the SimBA application to score cumulative time, frequency of entries, and latency to enter experimenter defined regions-of-interest within mazes. Scores from the social recognition tests were based on the position of the mouse’s snout.

### 2.8 | Quantification and statistical analysis

All analyses were performed using R Statistical Software (v4.2.1; R Core Team 2022).

#### Aggression

Since some animals never engaged in aggression on any day of the test the data were “zero inflated”; to account for this we used a Tobit vector generalized linear model using the *vglm* function from the *VGAM* package (Yee, 2023) to test for significant effect of *Cross* (maternal, paternal), *Genotype* (WT, Het), and their interaction on offensive behavior duration: v*glm(OffensiveTime ∼ Cross + Genotype + Cross:Genotype*). Model contrasts were set to “contr.sum” for *Cross* and to “contr.treatment” for *Genotype* effects coding. A similar generalized linear model with the same independent variable terms and contrast effects coding was used to for significant effects on over-dispersed *OffensiveBout* count data using the *glm* function with a “quasipoisson” distribution family parameter. Male and female data for the above models were analyzed separately, as was data from each of three trials. Latency data was tested with the log-rank test and plotted with a survival curve, and animals that did not attack in a trial received “censored” values of 600s. Separate curves were generated for cross (maternal, paternal) and day, and we utilized the s*urvdiff* function from the *survival* package to test for effect of genotype (WT, Het) (Therneau and Grambsch, 2000; Therneau, 2023). We used the *lmer* function from the *lme4* package (Bates et al., 2015) to fit a linear mixed-model with a random-effect of animal *Subject,* to test for significant fixed-effects of *Cross* (maternal, paternal), *Genotype* (WT, Het), *Trial*, and their interactions on social behavior and non-social behavior durations: *lme4(BehaviorTime ∼ Cross * Genotype * Day + (1|Subject))*. Model contrasts were set to “contr.treatment” for *Genotype* and “contr.sum” for *Cross* and *Day* effects coding. Male and female data were analyzed separately. Analysis of Deviance tables with Type III Wald test statistics were calculated for each of the models described above using the *Anova* function within the *car* package (Fox and Weisberg, 2019) to determine which factors and their interactions significantly reduce deviance in the models. Post-hoc contrast analyses using the *emmeans* package (Lenth, 2023) determined whether maternal or paternal allele heterozygote behaviors were significantly different from WT littermates.

#### Social recognition

Tracking data from DeepLabCut and SimBA gave us nose-point data for time spent in regions of interest. Significant habituation (trials 2-9) and dishabituation (trial 10) in time spent investigating the cup containing the conspecific for each trial were fit to a linear mixed– model with a random-effect of animal *Subject* and a fixed-effect of trial using the *lmer* function from the *lme4* package (Bates et al., 2015): *lmer*(*Invest* ∼ *Trial* + *(1|Animal)).* All genotype by sex groups were analyzed in separate models. Custom contrast effects coding was used to compare Trial 1 to Trials 2 – 9, and Trial 9 to Trial 10.

#### Dominance

We calculated the mean wins per day for each mouse based on the total number of wins and total days of testing for its group. We used a binomial generalized linear model using the *glm* function in R to test for the effect of *Cross* (maternal, paternal), *Genotype* (WT, Het), and their interaction on rank (dominant, non-dominant) or average wins per day expressed as proportional data from 0 to 1 (i.e. avg. 0 wins/day = 0, and 3 wins/day = 1): *glm(Variable ∼ Cross + Genotype + Cross:Genotype)*. Contrast effects coding for both *Cross* and *Genotype* were set to “contr.sum”. Male and female data were analyzed separately. Analysis of Deviance tables with Type III Wald test statistics were then calculated using the *Anova* function within the *car* package (Fox and Weisberg, 2019) to determine which factors and their interactions significantly reduce deviance in the models.

#### Sociability and social novelty

Significant impacts of maternal or paternal allele *Ddc* mutations on behavior in the three-chambered box sociability and social novelty tasks were measured with linear mixed-models in R using the *lme4* package (Bates et al., 2015). The difference in cumulative duration (in seconds) of time spent in the empty compared to conspecific chamber, or investigating the conspecific or empty wire cup, were first fit to the following linear or generalized linear-mixed: *(g)lmer(Time ∼ Chamber + Cross + Genotype + Chamber:Cross + Chamber:Genotype + Cross:Genotype + Chamber:Cross:Genotype + (1|Subject)).* Normally distributed continuous data (i.e. time) used a gaussian family (link = **“identity”**), and count data used a quasipoisson family (link = **“log”**), distributions. In this model the main factors of *Chamber* (levels = conspecific, empty), *Cross* (levels = maternal, paternal) and *Genotype* (levels = WT, *Th*-Het, *Ddc*-Het, *Ddc;Th*-Het) are fixed effects, and *Subject* is a random intercept effect that accounts for each individual mouse subject scored simultaneously on both levels of chamber. Contrasts for *Chamber*, and *Cross* in the model were set to “contr.sum” and to “contr.treatment” for *Genotype* effects coding. Male and female data were analyzed separately. Analysis of Deviance tables with Type III Wald test statistics were then calculated using the *Anova* function within the *car* package (Fox and Weisberg, 2019) to determine which factors and their interactions significantly reduce deviance in the models. Significant *Chamber:Cross* interaction effects (treatment contrast) indicate social preference/novelty difference between wildtype controls of the two crosses. Significant *Chamber:Genotype* interactions indicate that a heterozygous mutation, regardless of parent-of-origin, impacts social preference or novelty phenotypes relative to their wild-type, same-sex littermates. Significant three-way interaction effects for *Chamber:Cross:Genotype* indicates that the parent-of-origin (i.e. putative imprinting effect) of the heterozygous mutation (*Th, Ddc, or ThDdc*) affects social preference or novelty relative to WT littermates. Post-hoc contrast analyses using the *emmeans* package (Lenth, 2023) determined whether maternal or paternal allele heterozygote behaviors were significantly different from WT littermates.

## 3 | Results

### 3.1 Paternal heterozygote *Ddc* males are quicker to attack in resident-intruder paradigm

We sought to identify how the maternal versus paternal *Ddc* alleles affect offspring aggression by using the resident-intruder paradigm to assess male aggressive behavior (Fig. 2A). We examined differences in duration, latency, and bouts both within and across trial days. Since each genotype group contained animals that did not attack over any days of the test, or only some days of the test, we utilized a survival analysis to examine the latency to attack. We observed that *Ddc*^+/-^ paternal-null allele heterozygotes exhibited a shorter latency to attack on Day 1 of the test compared to WT littermates (Fig. 2D, *p* = 0.01). *Ddc^+/-^* paternal-null allele heterozygotes also exhibited increased aggressive behavior on the first trial compared to WT littermates and maternal cross male offspring. We found that loss of the paternal allele affected the attack duration on Day 1 (Fig. 2B, Cross x Heterozygosity W(1) = 5.8, *p* = 0.016, paternal cross pairwise estimate *p* = 0.0047). Fisher’s exact test was used to determine that the association between genotype and the total number of days the animal was aggressive, and the result trended toward significance (Fig. 2C, FE *p* = 0.086).

**Fig 2.**
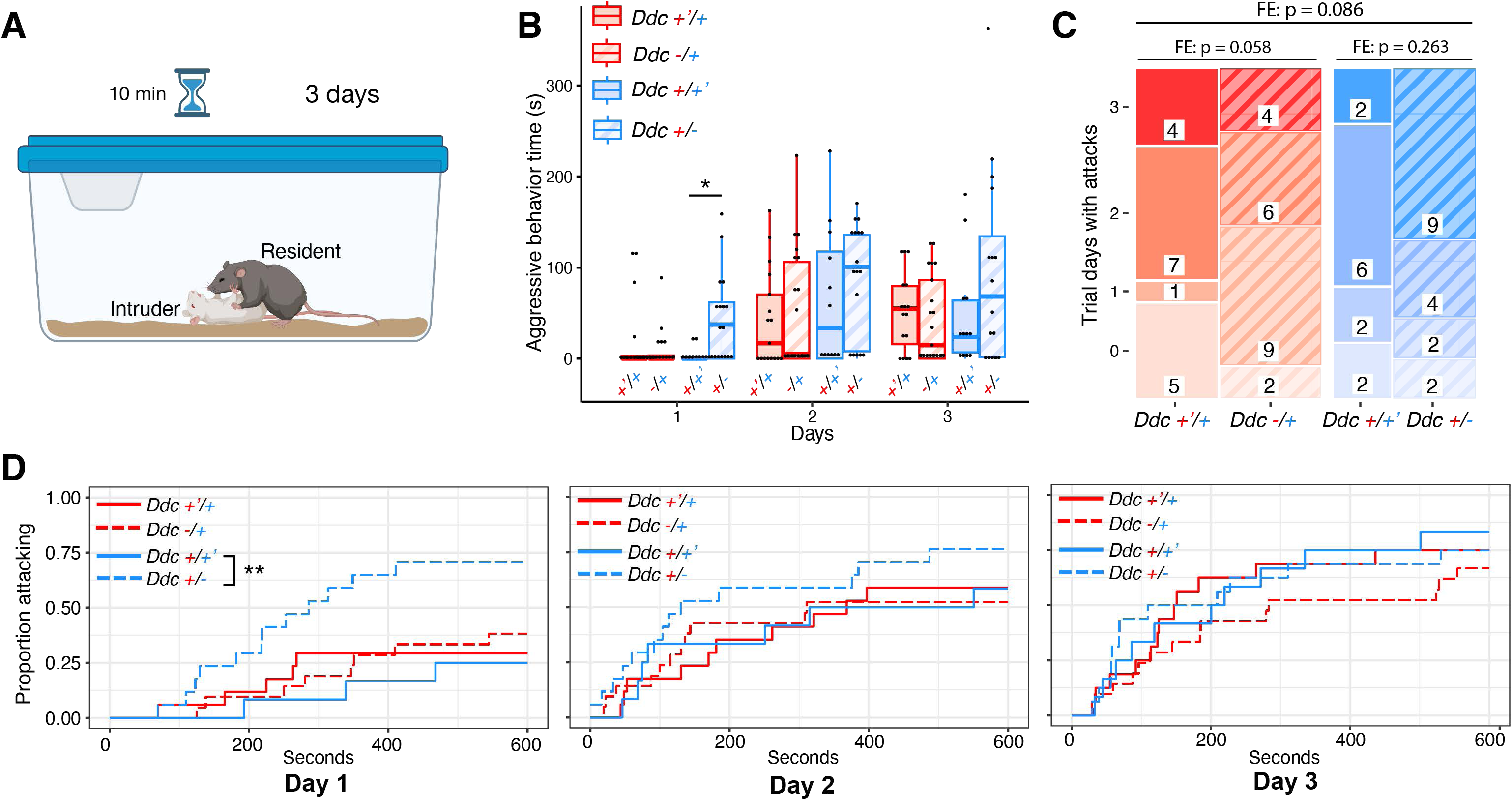
Paternal-null *Ddc* alleles affect aggressive behavior. **(a)** Testing occurred for 10 minutes a day over 3 days; an unfamiliar A/J strain mouse was placed in the subject’s home cage and aggression encounters, social exploration, and nonsocial behavior were scored. **(b)** Aggressive behavior duration (pushes, lunges, bites, and tumbling) shows paternal null-allele heterozygotes had significantly higher aggressive duration compared to their WT littermates on day 1, measured in seconds (s). **(c)** Comparison of number of days (n) attacking by genotype across all groups found a trend toward significance. **(d)** Survival plots show paternal heterozygotes are significantly faster to engage in aggressive behavior on Day 1 when compared to WT. Box plots indicate, median, interquartile range, and minimum/maximum range. Group genotypes: solid red *Ddc* +’/+, maternal WT; striped red *Ddc* -/+, maternal heterozygote; solid blue *Ddc* +/+’, paternal WT; striped blue *Ddc* +/-, paternal heterozygote. \**p* ≤ 0.05, \*\**p* ≤ 0.01.

Additional measures were consistent with increased aggression for *Ddc^+/-^* paternal-null allele heterozygotes. There was a significant interaction for the number of attack bouts on Day 1 (Fig. S1A, Cross x Heterozygosity W(1) = 4.2, *p* = 0.0396) and *Ddc^+/-^* paternal-null allele heterozygotes were significantly different from WT littermates (paternal cross pairwise estimate *p* = 0.013). For social exploration duration, an overall effect of day was observed (Fig. S1B, W(2) = 12.096, *p* = 0.002). For nonsocial behavior duration, an overall effect of heterozygosity was seen (Fig. S1C, W(1) = 4.6, *p* = 0.031), with no significant *Cross:Genotype* interaction indicative of parental allele specific (i.e. imprinting) effects.

### 3.2 *Ddc* paternal-null allele heterozygous males and maternal cross females have social recognition deficits

To determine whether *Ddc* imprinting affects social recognition, we measured interaction time during a ten-trial habituation/dishabituation task (Fig. 3A). For male *investigation zon*e time, each genotype group habituates to the social recognition task by Trial 5, with a significant difference in time spent investigating from Trial 1 (Fig. 3B, Table 4). For the dishabituation trial, *Ddc^-/+^*maternal-null allele heterozygotes had a significant difference in investigation time between Trials 9 and 10 (*p* = 0.007) and their WT littermates (*Ddc^+’/+^*) were trending towards a significant difference (*p* = 0.061). Meanwhile, *Ddc^+/-^* paternal-null allele heterozygotes did not have a significant difference in investigation time between trials 9 and 10 (*p* = 0.169), but their WT littermates did (*Ddc^+/+’^*, *p* = 0.003). For the *avoidance zone* (Fig. S2A, Table 5), only paternal WT (*Ddc^+/+’^*) showed a significant difference between Trials 9 and 10 (*p* = 0.003). Paternal-null heterozygotes (*Ddc^+/-^, p* = 0.238), maternal-null heterozygotes (*Ddc^-/+^, p* = 0.661), and maternal WT (*Ddc^+’/+^*, *p* = 0.121) did not show a difference in avoidance during the dishabituation trial.

**Fig 3.**
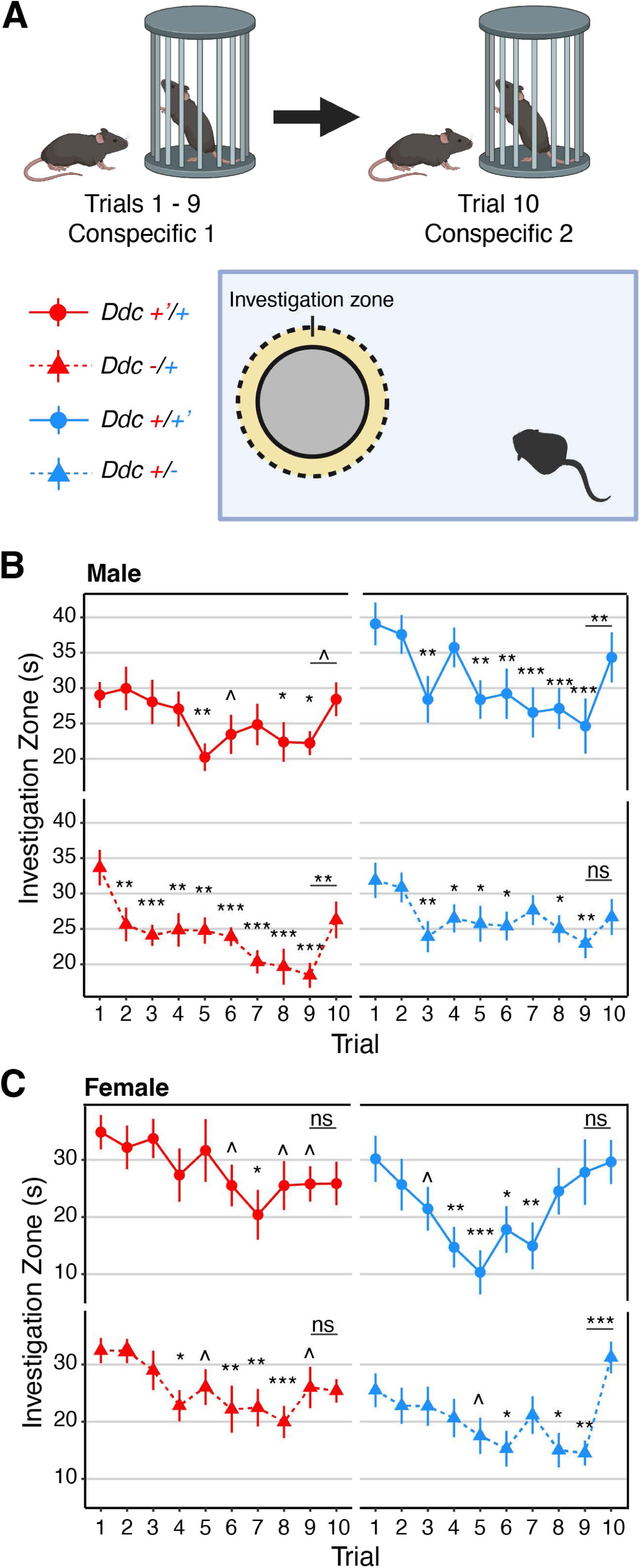
Inheritance of paternal-null mutant *Ddc* allele impairs social recognition in males. **(a)** Subject mice are scored for habituation of investigation of a confined same-sex conspecific for nine 1 min. trials with 10 min. inter-trial intervals. On the 10^th^ trial a new same-sex conspecific is used to measure social recognition by investigation dishabituation. **(b)** Nose point investigation, measured in seconds (s), within 2cm around cup of stimulus shows paternal-null allele heterozygous males do not dishabituate on the 10^th^ trial. **(c)** Investigation of conspecific by females shows only paternal-null allele heterozygotes dishabituate on the 10^th^ trial. Point-range plots indicated mean +/-se values for each of the following groups: solid red line, circle *Ddc* +’/+, maternal WT; dashed red line, triangle *Ddc* -/+, maternal heterozygote; solid blue line, circle *Ddc* +/+’, paternal WT; dashed blue line, triangle *Ddc* +/-, paternal heterozygote. *^p* ≤ 0.1, *\*p* ≤ 0.05, \*\**p* ≤ 0.01, \*\*\**p* ≤ 0.001.

**Table 4.**
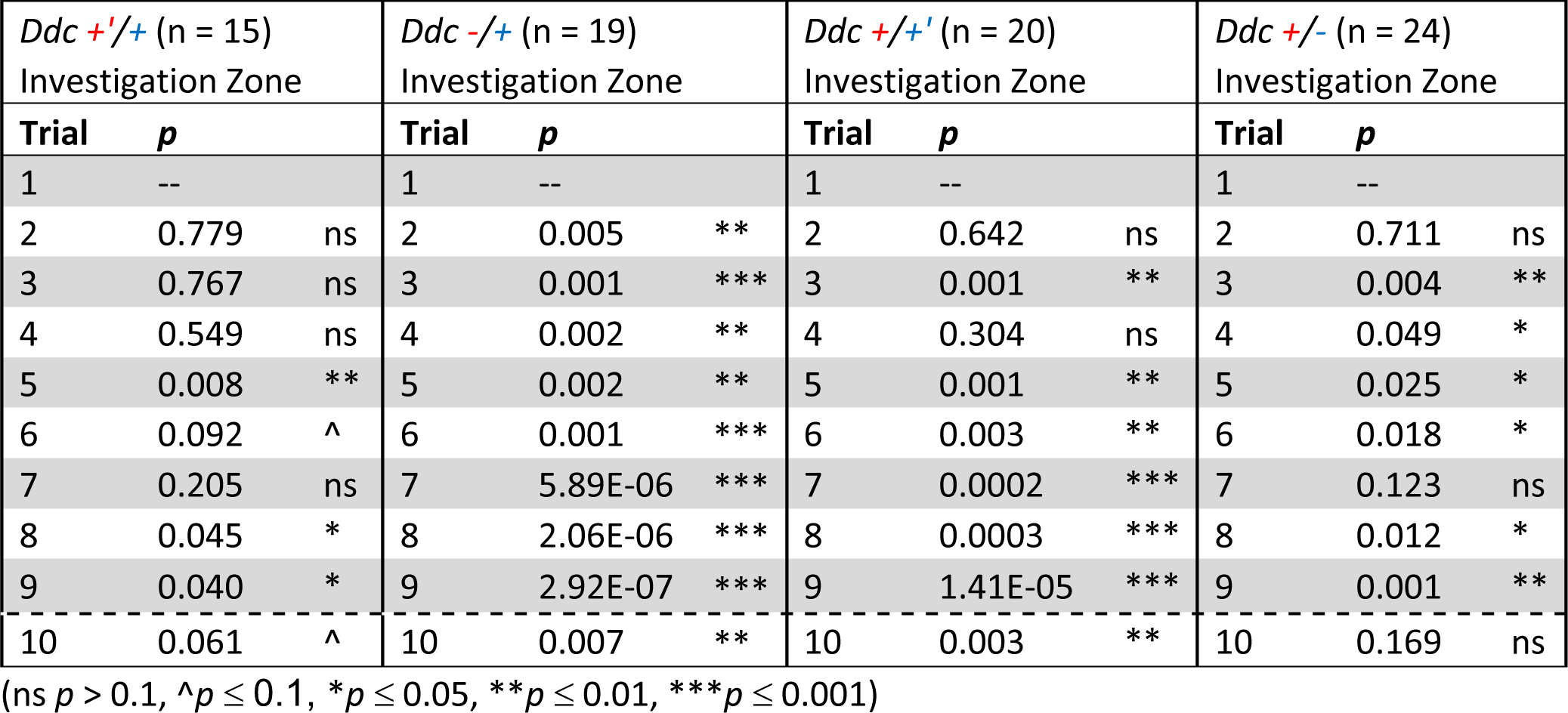
Male social recognition investigation zone *p* values by trial and genotype.

**Table 5.**
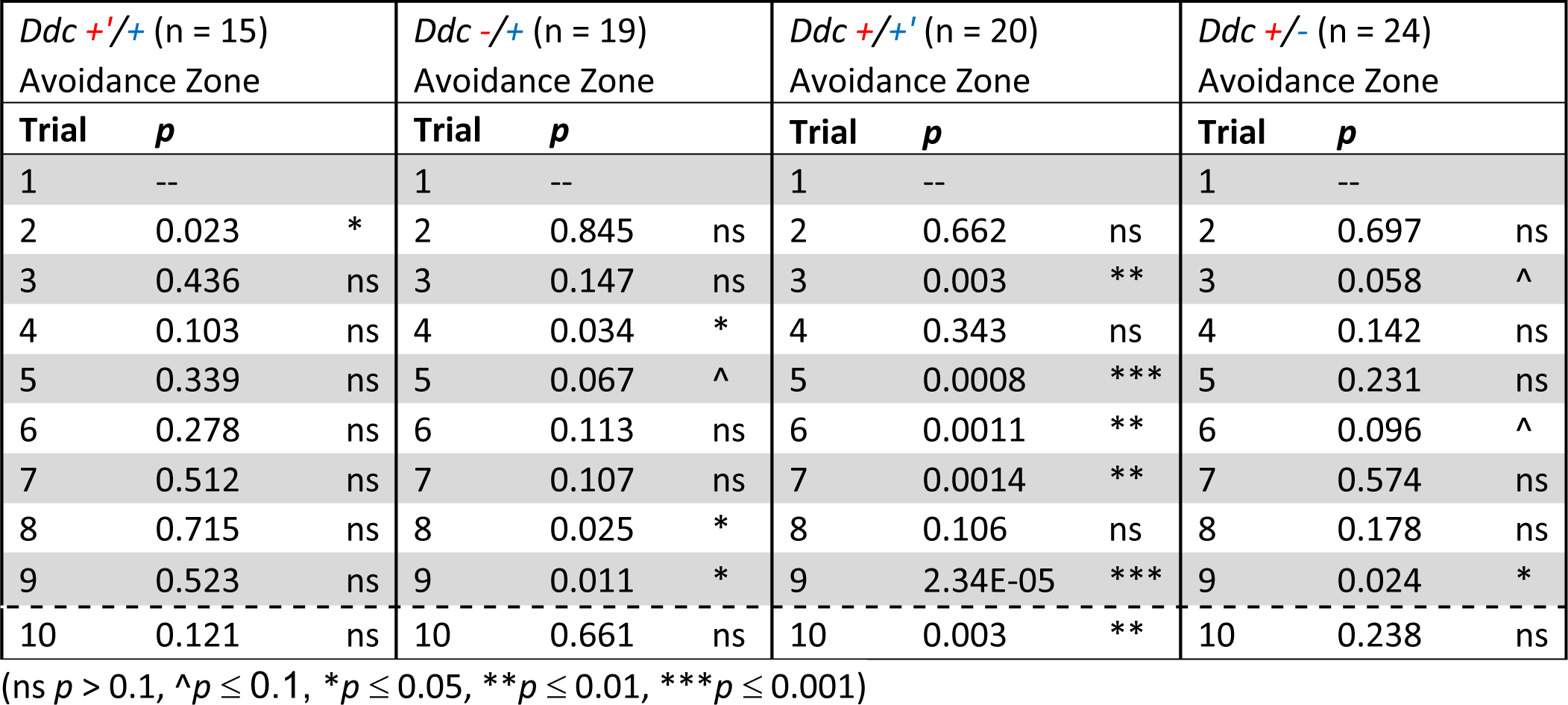
Male social recognition avoidance zone *p* values by trial and genotype.

For female *investigation zone* time, each genotype group appears to habituate to the task by Trial 6, with a significant difference in time spent investigating the conspecific from Trial 1 (Fig. 3C, Table 6). For the dishabituation trial, only *Ddc^+/-^* paternal-null allele heterozygotes had a significant difference in investigation between Trials 9 and 10 (*p* = 0.0001) and their WT littermates did not show a significant difference in dishabituation (*Ddc*^+/+’^, *p* = 0.705). For *Ddc* maternal cross female offspring, neither *Ddc^-/+^* maternal-null allele heterozygotes (*p* = 0.877) nor their WT littermates (*Ddc*^+’/+^, *p* = 0.990) showed a significant difference between Trial 9 and 10. Accordingly, a similar, but inverse, pattern is observed for time spent in the *avoidance zone* (Fig. S2B, Table 7). *Ddc^+/-^* paternal-null allele female heterozygotes had a significant difference in avoidance zone time between trial 9 and 10 (*p* = 0.0018), and their WT female littermates trended towards a significant difference (*Ddc*^+/+^’, *p* = 0.056). For the maternal cross females, neither heterozygotes (*Ddc^-/+^*, *p* = 0.522) nor their WT littermates (*Ddc^+’/+^, p* = 0.424) showed a significant difference between Trial 9 and 10.

**Table 6.**
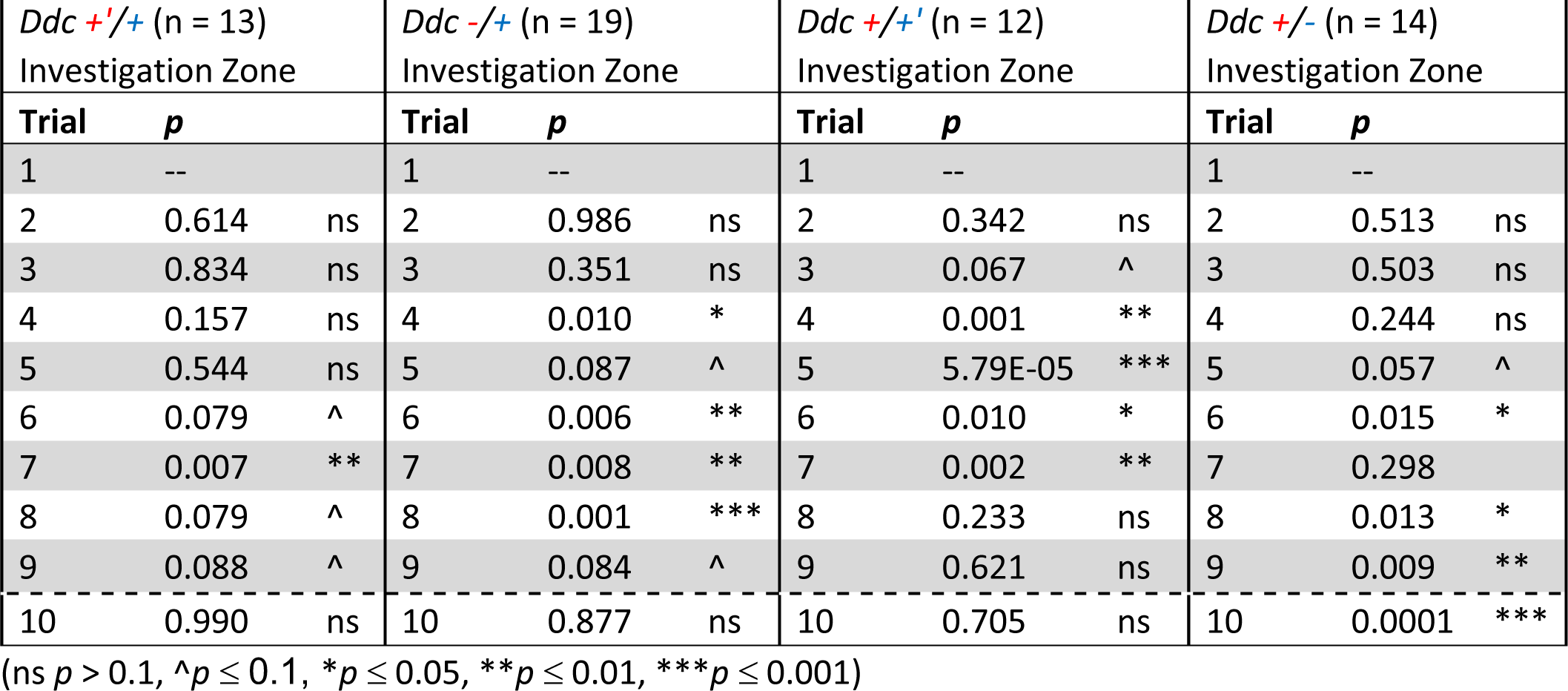
Female social recognition investigation zone *p* values by trial and genotype.

**Table 7.**
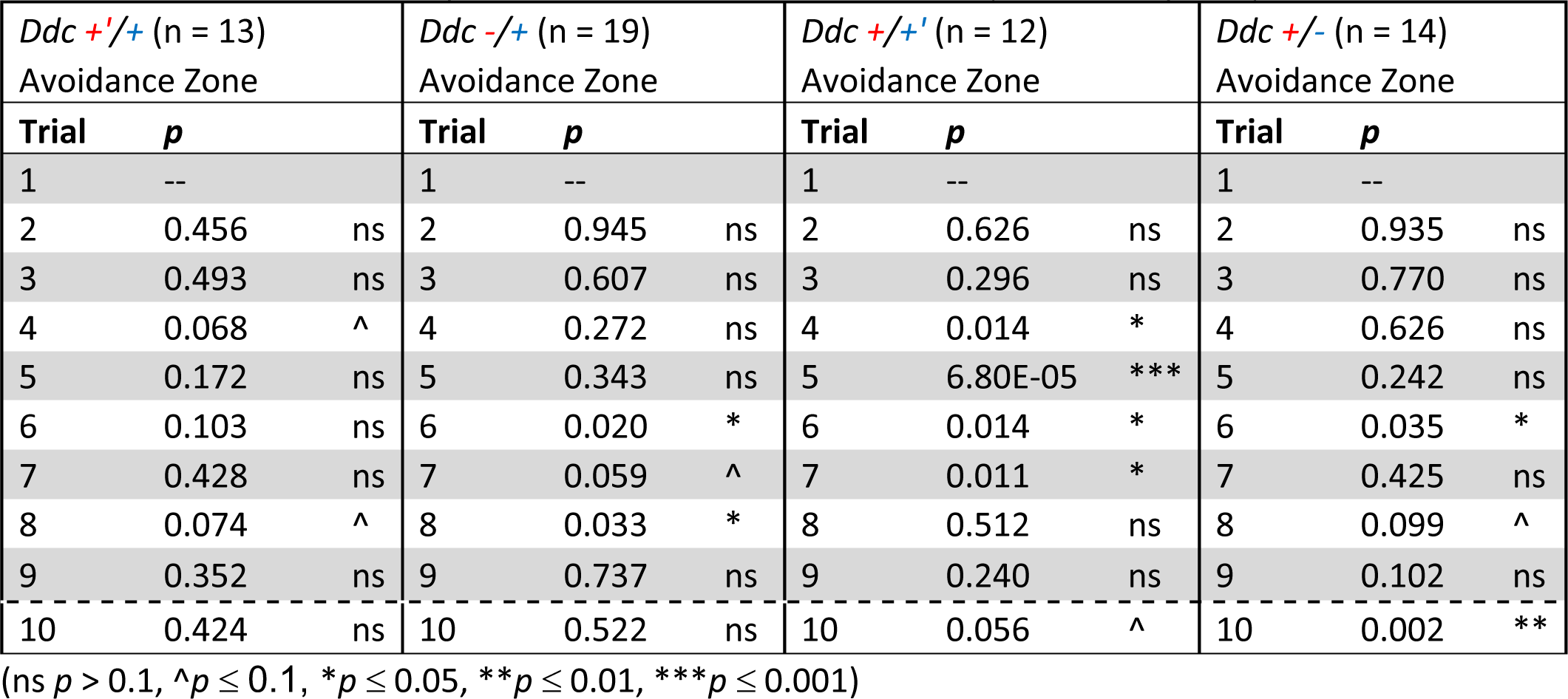
Female social recognition avoidance zone *p* values by trial and genotype.

### 3.3 *Ddc* maternal-null and paternal-null heterozygous mutations do not affect dominance

We utilized the tube test to assess social dominance phenotypes between co-housed reciprocal cross offspring. We housed four mice, one of each unique *Ddc* reciprocal cross genotype (Fig 4A), together starting when they were between P21 and P35 (Tables 2, 3) and waited until they were P60 to begin the tube test to assess established social dominance hierarchy. We found no significant effects of cross or heterozygosity on average wins per day in males (Fig. 4B, *p* = 0.404) or in females (Fig. 4C, *p* = 0.305). To assess differences in final rank, we looked at dominant animals against non-dominant animals (final rank of 2^nd^, 3^rd^, or 4^th^). We did not observe any significant differences between *Ddc* heterozygous mice and WT littermates in dominance ranking in males (Fig. 4D, FE *p* = 0.316) or in females (Fig. 4e, FE *p* = 0.316).

**Fig 4.**
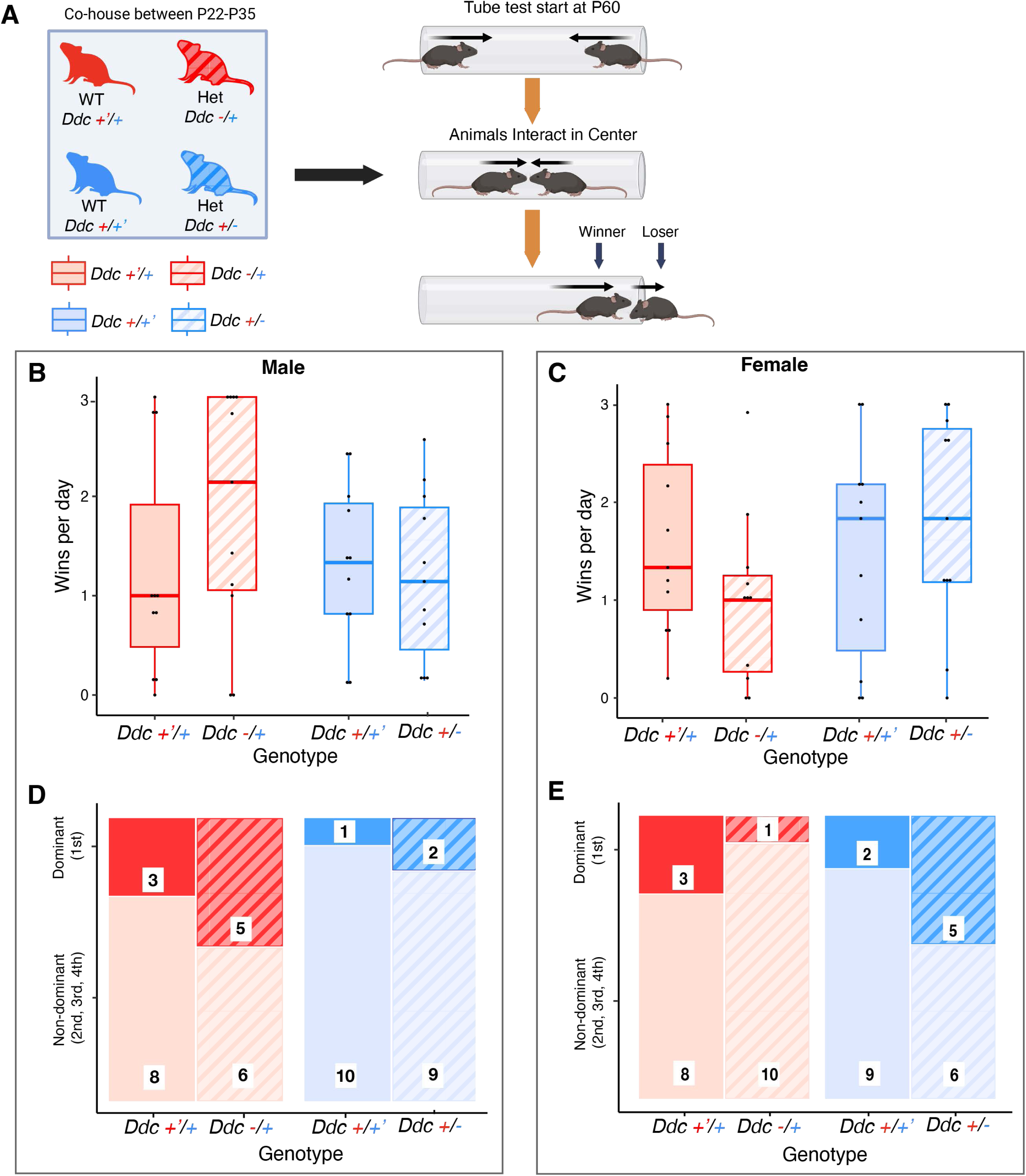
*Ddc* alleles do not affect dominance. **(a)** Age-matched, same-sex animals of each genotype were cohoused starting at between P22-P35 until testing as adults; at P60 tube test began to determine social hierarchy of cage mates. **(b)** No significant differences in average wins per day were observed in males or **(c)** females. **(d)** No significant differences in dominance rank were observed in males, **(e)** nor in females. Box plots indicate, median, interquartile range, and minimum/maximum range. Group genotypes: solid red *Ddc* +’/+, maternal WT; striped red *Ddc* -/+, maternal heterozygote; solid blue *Ddc* +/+’, paternal WT; striped blue *Ddc* +/-, paternal heterozygote.

### 3.4 Offspring from reciprocal compound heterozygote crosses show maternal *Th* allelic and *Ddc* allelic (both maternal and paternal) effects on sociability, and synergistic maternal *Th* and *Ddc* allelic effects on social novelty behaviors

Results from the *Ddc* heterozygous mutants found paternal allele effects on aggression and social recognition behaviors in males; however, noncanonical imprinting of both *Th* and *Ddc* in monoaminergic systems of the brain show maternal allele biased expression. Therefore, paternal allele effects on social behaviors were unexpected and not indicative of imprinted maternal allele expression in the brain regulating aggression and social recognition behaviors in males. To further examine potential functional effects of maternal allele imprinted expression in monoaminergic systems of the brain regulating social behaviors, we hypothesized that *Th* and *Ddc* maternal allele imprinted expression in the central nervous system may synergistically regulate social behavior functions. Thus, we tested offspring from reciprocal compound *Th* and *Ddc* heterozygous crosses (Fig. 5A) in sociability and social novelty tasks (Fig. 5B) to assess individual and synergistic catecholamine synthesis genes maternal and paternal allelic effects on these behaviors.

**Fig 5.**
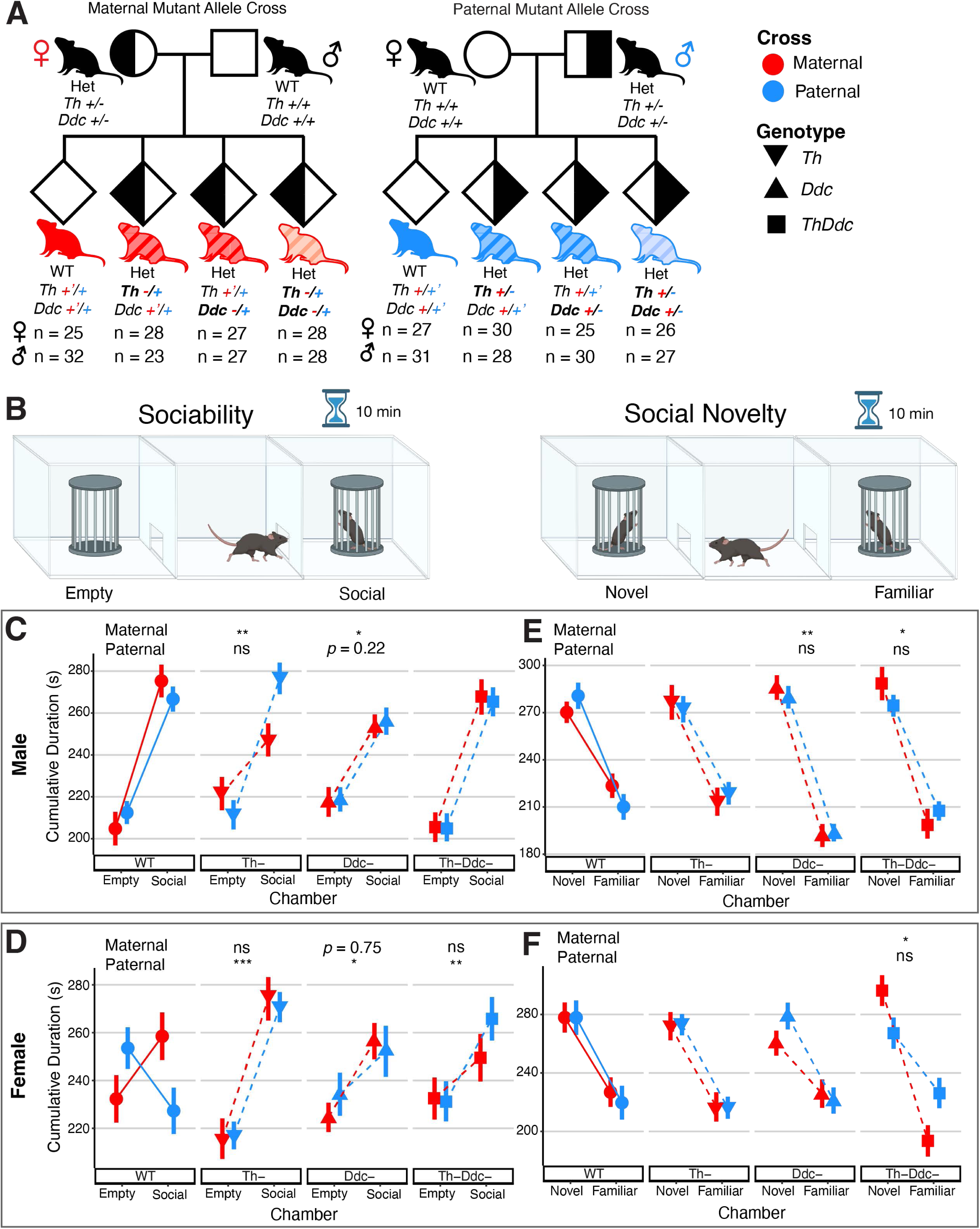
Sex specific *Th^-/+^*maternal-null allele, *Ddc* heterozygous null allele, and compound *Th^-/+^Ddc^-/+^*maternal-null allele effects on social preference phenotypes. **(a)** Reciprocal compound *ThDdc* maternal and paternal heterozygous mutant crosses result in WT, *Th* heterozygous, *Ddc* heterozygous, and *ThDdc* heterozygous offspring. **(b)** Subject mice were tested in two 10 min phases, first for sociability and then for social novelty. The sociability phase tested time spent investigating a cup containing a social stimulus or an empty cup. The social novelty phase tested time spent investigating the familiar stimulus from the first phase or a novel social stimulus mouse. **(c)** Male sociability results show *Ddc* heterozygotes, regardless of parental origin, and maternal-null *Th^-/+^* heterozygotes have a decreased preference for the social chamber compared to WT littermates, measured in seconds (s). **(d)** Female sociability results show a cross effect in WT offspring. **(d)** Male social novelty results show decreased preference for time in the chamber with the familiar conspecific for *Ddc* null allele heterozygotes (*Ddc^-/+^*and *Ddc^+/-^*) compared to WT littermates, and *Th^-/+^Ddc^-/+^* maternal-null allele heterozygotes when compared to maternal WT littermates. **(e)** Female social novelty results also show decreased preference for time in the chamber with the familiar conspecific for *Th^-/+^Ddc^-/+^* maternal-null allele heterozygotes compared to WT littermates. Point-range plots indicated mean +/-se values. Legend symbols indicating cross and *Th* and *Ddc* genotypes: blue, paternal cross; red, maternal cross; circles, WT; inverted triangles, *Th* heterozygotes; triangles, *Ddc* heterozygotes; squares, *ThDdc* compound heterozygotes. Following symbols indicate significant post-tests comparing heterozygotes of a particular cross to their respective WT littermates: \**p* ≤ 0.05, \*\**p* ≤ 0.01, \*\*\**p* ≤ 0.001.

A significant difference in the amount of total time spent in the chamber with a conspecific stimulus male compared to the empty chamber revealed, as expected, a strong preference for sociability in WT (*Th^+/+^;Ddc^+/+^*) males (Fig. 5c, *p* < 10^-16^). *Ddc* heterozygous males (*Ddc^-/+^* and *Ddc^+/-^*), regardless of whether the mutant allele was inherited from the mother or father, showed less preference to spend time in a chamber containing a male conspecific compared to WT littermates (Fig. 5C). A significant *Chamber:Genotype* interaction (*p <* 0.01) showed *Ddc^-/+^ and Ddc^+/-^* heterozygous males spent less time in the conspecific chamber and more time in the empty chamber compared to their WT littermates. The reduction in total-time sociability preference is a consequence of a significant reduction in the mean amount of time *Ddc* heterozygotes spent per visit in the chamber with the conspecific (Fig. S3A, *p* = 0.0186). Thus, *Ddc* heterozygotes were departing the chamber with the conspecific more quickly on average, and lingering less, than their WT littermates. These results are indicative of a *heterozygous* allele effect on sociability.

In contrast to the heterozygous effects of *Ddc*, a significant interaction between *Cross* (parent-of-origin of mutant allele) and inheritance of a *Th* heterozygous allele was found for cumulative chamber-time sociability with a conspecific (Fig. 5C, *p* = 0.005) during the sociability task. Post-test contrasts revealed that males inheriting a mutant maternal *Th* allele had significantly less preference for the conspecific relative to the empty chamber compared to WT littermates (Fig. 5C, *p* = 0.004). However, males with a paternal *Th* mutation were no different from WT. Per visit, *Th^-/+^* maternal-null allele heterozygotes spent no more time in the conspecific chamber than the empty chamber before departing (Fig. S3A, *p* = 0.008), whereas paternal heterozygotes and WT littermates of both crosses remained in the conspecific chamber longer than the empty chamber before leaving. In addition, a significant interaction of *Cross* and *Th* mutation (*p* = 0.007) and post-tests revealed that *Th* maternal-null allele heterozygotes (Fig. S3B, *p* = 0.002), but not paternal-null allele heterozygotes, had a reduced preference for investigating (i.e. sniffing) the cup containing the conspecific than the empty cup compared to WT littermates. Further data indicate that the reduction in conspecific chamber/cup preference results in *Th* maternal-null allele heterozygote males spending more time in the center chamber of the three-chambered box (Fig. S3C, *p* = 0.053).

A significant effect of *Chamber* (total time) by *Cross* effect (Fig. 5D, *p* = 0.003) showed that WT female offspring from the paternal cross had an aversion to sociability with a conspecific male compared to a sociability preference for WT female offspring from the maternal cross. Significant interactions between *Chamber* preference and *Genotype* (*Ddc*, *p* = 0.046); and *Chamber* preference by *Cross* by *Genotype* (*Th, p* = 0.0498; *ThDdc*, *p* = 0.0048) interactions and post-tests (*Th^+/-^, p* < 0.0001; *Th^+/-^Ddc^+/-^*, *p* = 0.0012) showed that the sociability aversion in WT females from the paternal cross is eliminated in their mutant sister littermates (Fig. 5D). Further analysis of the data shows that the social aversion in WT females from the paternal cross is driven by a reduction in the amount of time they spend per visit in the chamber with the conspecific male compared to the empty chamber (Fig. S4A, *p* = 0.022) and not by reductions in the frequency (n) or latency (s) to visit the conspecific relative to the empty chamber *(data not shown).* A significant *Cup* investigation (total time investigating) by *Genotype* interaction (Fig. S4B, *p* = 0.007) revealed that *Th* heterozygosity increased preference to investigate a conspecific. Finally, an interaction of *Cross* by *Genotype* (Fig. S4C, *p* = 0.005) showed that in females paternal inheritance of compound *Th^+/-^Ddc^+/-^*mutations reduced time in center relative to WT littermates. Overall, WT females from mutant fathers showed a lack of a sociability relative to WT females from mutant mothers and heterozygous mutants regardless of parental origin of the mutant *Th* and *Ddc* alleles.

After testing animals for social preference, animals were returned to the center of the three-chambered box, and a new unfamiliar conspecific was placed into the previously empty cup before allowing the mice to freely explore. As expected, WT males had a strong preference for total time (s) in the novel stimulus mouse chamber (Fig. 5e, *p* < 10^-13^) and for exploring the cup with the “novel” compared to the now “familiar” conspecific (Fig. S5B, *p* < 10^-11^). A significant interaction for time in *Chamber* by *Genotype* (Fig. 5E, *p* = 0.0045) indicates an increased preference for the novel conspecific in *Ddc* heterozygotes that inherit mutant alleles from either their mother or father. A chamber by genotype by cross three-way interaction (*p* = 0.0385) and post-tests indicated that male *Th^-/+^Ddc^-/+^* compound heterozygotes that inherit mutant alleles from their mothers but not their fathers have an increased preference for a novel conspecific relative to their WT littermates (Fig. 5E, *p* = 0.014). A *Chamber* by *Genotype* by *Cross* 3-way interaction (Fig. S5A, *p* = 0.022) indicates that a preference for frequency (number) of visits to the chamber with the novel mouse in male WT offspring is lost in *Th^+/-^* heterozygotes that inherit a paternal-null allele (Fig. S5A, *p* = 0.011). A significant *Cup* by *Genotype* interaction (Fig. S5B, *p* = 0.041) shows increased preference for investigating the cup containing the novel mouse over the familiar compared to WT in both maternally and paternally inherited *Ddc* heterozygotes, while a significant *Cup* by *Genotype* by *Cross* interaction (Fig. S5B, *p* = 0.039) and post-tests show the enhanced novel cup preference is only significant for maternally inherited compound *Th^-/+^Ddc^-/+^* male heterozygotes (*p* = 0.012). Additional measures for males show that *Ddc* heterozygous mutations (maternal and paternally inherited) increases the time in (Fig. S5C, *p* = 0.0008), and frequency of visits to (Fig. S5D, *p* = 0.038), the center chamber. Male *Th* heterozygotes from both maternal and paternal crosses less frequently enter the center chamber (Fig. S5D, *p* = 0.005).

Like WT males, WT females have a strong preference for spending time in the chamber with a novel mouse compared to the familiar conspecific (Fig. 5F, *p* < 10^-7^). Neither a maternal nor paternal allele inherited heterozygous mutation of either *Th* or *Ddc* alone influences this preference for social novelty (Fig. 5F). However, a significant interaction of *Chamber* by *Cross* by *Genotype* effect (Fig. 5F, *p* = 0.015) and post-tests reveal that, when compared to WT littermates, females inheriting both *Th^-/+^* and *Ddc^-/+^* maternal-null heterozygous alleles have a significant increase of time in the chamber with the novel conspecific (Fig. 5F, *p* = 0.02). Compound *Th^+/-^ Ddc^+/-^* paternal-null allele heterozygotes have no difference from WT littermates in preference for time in the novel conspecific chamber; however, a significant effect of *Genotype* shows that paternal *Th^+/-^Ddc^+/-^*heterozygotes spend more time investigating the cups of both familiar and novel conspecifics compared with WT littermates with no increased preference for the novel conspecific (Fig. S6A, *p* = 0.018). Female *Th* heterozygotes, regardless of parental origin of the null allele, spend more time in (Fig. S6B, *p* = 0.023) and more frequently visit (Fig. S6C, *p* = 0.03) the center of the three-chambered box. Similarly, *Ddc* heterozygotes, regardless of parent of origin, show an increased locomotor behavior by more frequently visiting the center (Fig. S6C, *p* = 0.005). Overall, these results are consistent with maternal biased expression of these two enzymes in the catecholamine synthesis pathway synergistically regulating offspring social novelty preferences.

## 4 | Discussion

Disruptions of genomic imprinting can cause genetic disorders with mental deficiencies that include atypical social behavior (Isles et al., 2006; Wilkinson et al., 2007). We hypothesized that *noncanonically* imprinted allelic expression effects of monoaminergic genes may functionally regulate social behaviors. In *Th^-/+^* and/or *Ddc^-/+^* maternal-null heterozygotes, imprinted brain cell types that normally have maternal monoallelic or maternal allele dominant expression of *Th* and/or *Ddc*, should have absent or greatly reduced functional capacity to synthesize and release monoamine neurotransmitters (Fig. 1). Conversely, imprinted cell types with paternal monoallelic or paternal dominant allelic expression (i.e. adrenal chromaffin cells) should have absent or diminished capacity to synthesize and release monoamines in *Th^+/-^* and/or *Ddc^+/-^* paternal-null heterozygotes. Here we show that males with *Ddc^+/-^* paternal-null heterozygous mutations displayed altered aggression and social recognition, but not dominance behavior. We also found that males with *Th^-/+^* maternal-null heterozygous mutations have decreased preference for social interaction during a sociability test, and compound *Th^-/+^Ddc^-/+^* maternal-null heterozygotes show an overall increased preference for novel over familiar conspecifics. These findings are consistent with differential functional roles for imprinted expression of maternal *Th* alleles and paternal *Ddc* alleles in the regulation of discrete social behaviors.

We found males with *Ddc^+/-^* paternal-null heterozygous alleles were significantly more aggressive on the first day of the test than paternal cross WT (*Ddc^+/+’^)* littermates, and WT and mutant maternal cross males (*Ddc^+’/+^* and *Ddc^-^*^/+^; Fig. 2B). This was a surprising finding since *Ddc*^+/-^ paternal-null heterozygotes are expected to have a loss of function for paternal-allele biased expressing cells while *Ddc*^-/+^ maternal-null heterozygotes are expected to have a loss of function for maternal-biased cells in the brain. Peripheral catecholamines are primarily secreted from the adrenal gland, where there is *Ddc* paternal-allele biased expression (Babak et al., 2015; Bonthuis et al., 2022), have been shown to play roles in arousal prior to, and engagement in, aggression; however, the focus on behavioral mechanisms has largely been on central systems (Sgoifo et al., 1996; Haller et al., 1997; Pohorecky et al., 2004). Two monoamine neurotransmitters, 5-HT and DA have been routinely linked to aggression in many species (Almeida et al., 2005; Nelson and Trainor, 2007; Seo et al., 2008). Low 5-HT levels are associated with increased impulsive and aggressive behavior in mice (Serri and Ely, 1984; Miczek et al., 2001; Mosienko et al., 2012) and 5-HT neurons in the DR of rats show increased *c-fos* expression following aggressive encounters (Vegt et al., 2003). DA release from the nucleus accumbens (NAc) has been observed following aggressive interactions in rats, and optogenetic activation of DAergic neurons in the VTA of mice has been shown to increase aggression (Erp and Miczek, 2000; Yu et al., 2014). Aggressive behavior has been linked to other catecholamine synthesis genes. Mice null for *Dbh*, a synthesis gene in the pathway for NE and E, were shown to have both altered social memory and a reduction in aggression (Marino et al., 2005); and mice with over expression of *Pnmt*, encoding the synthesis enzyme for E, displayed heightened aggression (Sørensen et al., 2005).

Complex neural circuits of aggression receive inputs from the main and accessory olfactory bulbs that are sent to the medial amygdala (MEA) which then sends signals to several regions including the lateral septum (LS), bed nucleus of the stria terminalis (BNST), periaqueductal grey (PAG) and multiple regions of the hypothalamus including the medial preoptic area (mPOA), ventral premammillary nucleus (PMv), and ventromedial hypothalamus (VMH) (Nelson and Trainor, 2007; Tachikawa et al., 2013; Falkner et al., 2016; Pardo-Bellver et al., 2017; Chen and Hong, 2018; Wei et al., 2021). Interestingly, many regions implicated in aggression circuitry, including the BNST, PAG, mPOA, and PMy, were also found to contain cells with imprinted *Ddc* monoallelic expression of the maternal allele (Bonthuis et al., 2022). Aggressive behavior circuitry could be impacted by imprinting effects in multiple regions to regulate aggressive behaviors. The maternally expressed imprinted gene *Cdkn1c* is responsible for midbrain DAergic cell proliferation, and mouse models mimicking a loss of *Cdkn1c* imprinting have been found to exhibit increased aggression and unstable social hierarchy (McNamara et al., 2018b, 2018c).

Although aggression and dominance are often thought of as dependent, we did not observe *Ddc* maternal or paternal allelic effects on social hierarchy and dominance. Dominant animals often use aggressive interactions to establish hierarchy, but top-ranking animals are not necessarily the most aggressive (Wang et al., 2014; Holekamp and Strauss, 2016). Social hierarchies are dependent on the ability to recognize a conspecific and their social status (Wang et al., 2014) and we show a deficit in social recognition in males with a *Ddc^+/-^* paternal-null heterozygous alleles. Though the animals eventually habituated to the social stimulus they did not dishabituate when a novel conspecific was introduced in the last trial of the task (Fig. 3B). Social recognition is thought to be strongly regulated by oxytocin (OT) and vasopressin (AVP) (Ferguson et al., 2002; Lee et al., 2008) but it has been shown that NE cell populations in the olfactory bulb are critical for OT and AVP binding for successful social recognition behavior (Dluzen et al., 1998). One aspect of social recognition is the ability to recognize kin and it has been proposed that imprinted genes might regulate this aspect of social recognition (Isles et al., 2006). One study utilizing offspring from reciprocal F1 hybrid mice from CBA/Ca and C57Bl/6 strains found both males and females were avoidant of female urinary odors from mice of the same strain as their mothers (Isles et al., 2001). Interestingly, only our female *Ddc^+/-^* paternal-null heterozygotes showed significant dishabituation in the social recognition task (Fig. 3C). The lack of dishabituation in both maternal and paternal WT females was particularly surprising, however this could be due to several reasons. Female rats have been shown to demonstrate less initial social investigation than males but more sustained social memory (Markham and Juraska, 2007) and in a habituation/dishabituation task male mice investigate both the familiar and novel stimulus more overall than females (Tejada and Rissman, 2012). Hormones have also been shown to play a role in social recognition – in female mice, improved social recognition has been associated with the high levels of circulating estrogens and progesterone during the proestrus phase of the estrous cycle (Aspesi and Choleris, 2022). A limitation of our study is that we did not account for estrous cycle in our female mice. Social recognition is also important in the decision making for aggressive behavior (Gabor et al., 2012). Many factors about the conspecific need to be processed, including their familiarity, reproductive status, and relatedness. Social recognition enables the individual to recall previous interactions and the emotional valence linked to those interactions (Gabor et al., 2012; Chiang et al., 2018). Having impaired social recognition could make any conspecific appear to be a threat regardless of the number of times they have encountered each other, thus lowering the threshold to engage in aggression. Importantly, our studies found that paternally inherited *Ddc* alleles, and not maternally inherited alleles, regulate male aggression and social recognition phenotypes, which is inconsistent with the observed maternal allele biased *Ddc* expression in the adult brain functionally regulating these behaviors, but is consistent with functional regulation by *Ddc* paternal allele expression bias in the periphery (Bonthuis et al., 2022).

*Ddc* is not the only imprinted gene in the catecholamine synthesis pathway; *Th* also exhibits a maternal allele bias in the brain (Bonthuis et al., 2015). It could be that noncanonically imprinted maternal allele biased expression for both *Ddc* and *Th* could potentiate maternal imprinting functional effects in catecholaminergic cells which has been demonstrating for some aspects of foraging behavior (Bonthuis et al., 2022). The sociability/social novelty paradigm provides insights into a few aspects of social behavior and serves as a mouse model for autism symptoms. The first phase, sociability, gives us a look at social approach and preference, while the second phase, allows for evaluation of social novelty discrimination and social memory (Moy et al., 2004; Silverman et al., 2010). Male heterozygous *Ddc* null allele offspring from reciprocal compound *ThDdc* heterozygous crosses spend less time in the chamber with a conspecific regardless of parent-of-origin of the mutation (Fig. 5C). The upshot of this result is that expression from both *Ddc* alleles (*Ddc^+/+^*) is required for normal WT social preference but is not indicative of *Ddc* genomic imprinting effects regulating sociability. This *heterozygous-null* mutation effect on sociability is somewhat surprising since *Ddc* is not considered to be a rate limiting enzyme in monoamine synthesis pathways, and as such, gene dose in biallelic expressing (non-imprinted) cells would not be predicted to affect overall neurotransmitter levels (Flatmark, 2000; Kaushik et al., 2007; Daubner et al., 2011). Male *Th*^-/+^ maternal-null allele heterozygotes also show a decreased preference for investigating a conspecific compared to their WT littermates during the sociability test (Fig. 5C). A closer look at the data revealed *Th^-/+^* maternal-null heterozygotes spend on average the same amount of time per visit to both the social chamber and the empty chamber, while their WT littermates spend longer visits to the social chamber (Fig. S3A). The *Th^-/+^* maternal-null males also spend more time in the center chamber than their WT littermates (Fig S3C). Together, these results are consistent with the hypothesis that cells in the brain (or elsewhere) with maternally biased *Th* expression (i.e. *Th* imprinted cells) impact the function of circuits (or other systems) regulating sociability behaviors when inheriting a maternal mutant allele, whereas inheritance of a paternal allele mutation does not.

The social novelty phase examines the mouse’s natural preference to investigate a new social stimulus as well as to recognize and remember the difference between the familiar and novel stimuli (Moy et al., 2004; Silverman et al., 2010). We also saw *Ddc* heterozygous-null effects on social novelty regardless of null allele parent-of-origin, with both maternal and paternal heterozygote males showing decreased interest in the familiar conspecific (Fig. 5E). Both male and female compound *Th^-/+^Ddc^-/+^* maternal-null allele heterozygotes showed an increased preference for the novel conspecific relative to the familiar conspecific compared to their WT littermates (Fig. 5E, F). These results indicate maternal allele imprinted expression of two genes in the catecholamine synthesis pathway synergistically regulate social novelty preference, and support our other findings that imprinted allele specific expression effects can influence behavior.

As stated above, we found behavioral effects on aggression and social recognition in our *Ddc^+/-^*paternal-null allele heterozygous males. We previously reported noncanonical imprinting with *Ddc* maternal allele biased expression in the brain, and *Ddc* paternal allele biased expression in the adrenal medulla (Bonthuis et al., 2022). These previous findings indicated that genomic imprinting effects on *Ddc* allelic expression is highly cell-type specific with maternal monoallelic expressing subpopulations centrally (i.e., adult brain) and predominant paternal monoallelic expressing subpopulations (i.e. adult adrenal medulla, and developing heart) peripherally (Babak et al., 2015; Bonthuis et al., 2022; Juan et al., 2022). Cell-and tissue-type specific imprinting has been reported for other genes and there are a handful of imprinted genes which have been reported to change expression of their parentally inherited alleles during development (Laukoter et al., 2020). Laukoter and colleagues (2020) sought to analyze cell-type specific imprinting in the cerebral cortex for 25 known imprinted genes at several timepoints between embryonic day 15 (E15) to P42, and found that *Snrpn*, *Ndn*, and *Cdkn1c*, showed marked cell-type-specific expression changes during development. *Grb10* is another imprinted gene which switches from maternal to paternal allele biased expression during development, changing in a region dependent manner from maternal biased expression at E15 in whole-brain tissue to paternal biased expression before P0 in the hypothalamus, before P8 in the cerebral cortex, and before P15 in the cerebellum (Perez et al., 2015). *Grb10*, which is in the same imprinted gene cluster as *Ddc*, continues to be expressed from the paternal allele in adult monoaminergic brain regions and maternal allele in most peripheral tissues; in contrast, *Ddc* exhibits maternal allele biased expression in the adult brain and paternal allele expression in the periphery (adult adrenal gland, embryonic heart) (Garfield et al., 2011; Plasschaert and Bartolomei, 2014; Sheng et al., 2022). In adult mice, the *Grb10* paternal allele has been shown to be involved in the regulation of behavior, including dominance, stability of social hierarchies, and risk-taking behavior (Garfield et al., 2011; Rienecker et al., 2019; Dent et al., 2020). *Grb10* and *Ddc* imprinted expression was recently found to be epigenetically regulated by a secondary differentially methylated region (DMR) that is necessary for tissue-specific, monoallelic expression in the embryonic heart (Juan et al., 2022). It is possible that like *Grb10* imprinted expression in the brain, *Ddc* also undergoes similar epigenetic allelic expression switches during development, and loss of hypothetical paternal allele expression in *Ddc*^+/-^ paternal-null heterozygous mice during brain development could alter the function of adult brain circuitry (Perez et al., 2015). It has been shown that critical periods of neural development are sensitive to monoamine signals that impact sensory system development, anxiety and depression related behaviors, and aggression and impulsivity (Suri et al., 2015). Imprinted allelic expression effects on brain development vs. peripheral systems would need to be confirmed with maternal and paternal reciprocal central (CNS) vs. peripheral (PNS) nervous system, and temporal, conditional heterozygous mutation strategies.

These findings provide insights into potential social behavior functional consequences of cell type specific complex genomic architectures epigenetically regulating imprinted allelic expression in central and peripheral monoaminergic systems. We show that maternal and paternal alleles have differential effects on discrete social behaviors, but also heterozygous effects independent of the parent of origin implicate *Ddc* gene dosage effects on social behavior. Maternally inherited *Th* alleles impact aspects of social approach and preference for time with a conspecific. In compound *ThDdc* mice, maternally inherited *Th* and *Ddc* alleles seem to work synergistically to regulate preference for a novel social interaction over a familiar social stimulus, thus implicating functional roles for maternal biased allelic expression in the adult brain. On the other hand, *Ddc* paternal alleles influence aggression and social recognition in males, implicating *Ddc* paternal allele biased expression in the adult periphery, or during brain development, functionally affecting these behaviors. Heterozygous *Th* and/or *Ddc* null allele effects on the quality of parental care given by reciprocal cross moms and dads cannot be excluded for roles in reciprocal heterozygous offspring phenotypes. For example, allelic mutations in reciprocal cross mothers compared to fathers (Figs. 1B, 5A), could cause differences in pup environmental experiences either *in utero* or during postnatal parental care. However, we expect parental care influences on offspring phenotypes to equivalently affect WT littermates and is accounted for in our controlled experimental design and statistical analyses. Additional experimental strategies are needed to elucidate whether maternal and paternal allele specific influences on social behaviors are caused by noncanonical monoaminergic imprinting effects on allelic expression in the CNS or PNS at particular stages of development. Experimental evidence suggests that *DDC* may also exhibit imprinted allelic expression effects in the human brain (Babak et al., 2015), and future investigations could examine potential roles in human behavior. Elucidating allelic expression mechanisms that contribute to the function of central and peripheral monoaminergic systems and the heritability of complex behaviors may provide important insight into complex genetic risk factors underlying mental health disorders known to be affected by monoamine neurotransmitters (Gainetdinov and Caron, 2003).

In summary, this study shows that heterozygous allelic variants in monoaminergic systems can have functional consequences on social behaviors that depend on the parent-of-origin of their inheritance. The findings implicate significant roles for epigenetic allele-specific expression programs in the complex heritability and heterogeneity of social behaviors in systems also relevant to human mental health disorders.

## Acknowledgements

This work was supported by the National Institute of Mental Health; NIH grant K99/R00MH111912. We also thank Susan Steinwand and Dr. Noelle James for technical support, and Dr. Richard Palmiter for the Th-delta mouse line. Some of the images in the figures were created with BioRender.

## Author contributions

PJB: Conceptualization (lead); methodology; formal analysis; writing – review & editing; visualization; funding acquisition

MD: Investigation

RLE: Investigation

TH: Investigation

DDM: Resources

EMO: Conceptualization (supporting); methodology; investigation; formal analysis; writing – original draft; visualization

SJR: Software ALT: Software

## Author approval

All authors have seen and approved the manuscript; and it has not been accepted or published elsewhere.

---

**Figure S1.**
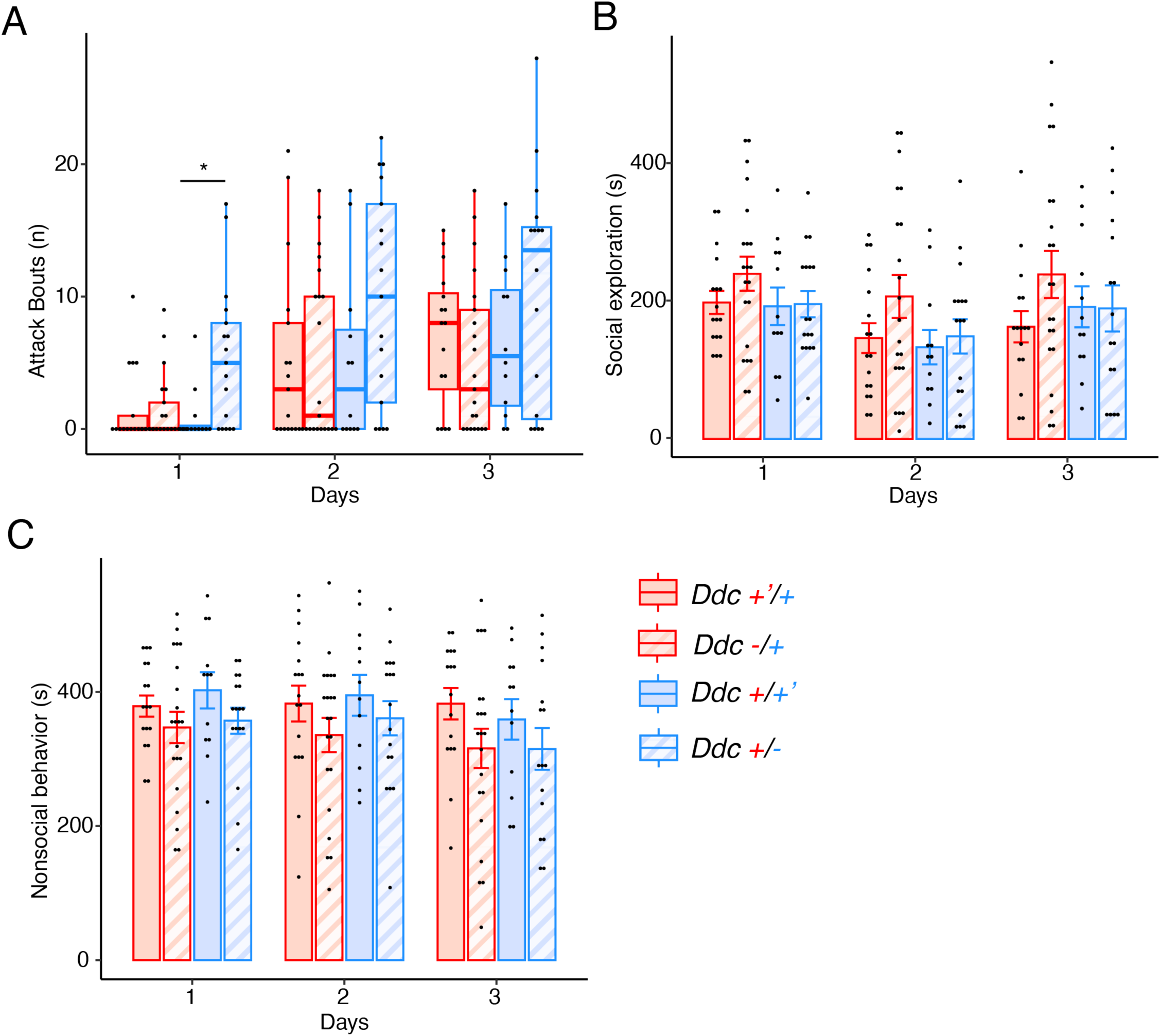
*Ddc+/-* paternal null allele heterozygotes have more offensive bouts on the first day of test, but no imprinting effects for social exploration or nonsocial behavior were observed. **(a)** There was a significant interaction for the number (n) of aggressive bouts on Day 1 and *Ddc^+/-^* paternal-null allele heterozygotes were significantly different from WT littermates. **(b)** Time spent in seconds (s) social investigating A/J intruder by genotype and day; an overall effect of day was observed but no effects of cross or heterozygosity were observed. **(c)** Nonsocial behavior during resident-intruder test for aggression in seconds (s); an overall effect of heterozygosity was seen, with no significant interaction indicative of parental allele specific (i.e. imprinting) effects. Box plots indicate, median, interquartile range, and minimum/maximum range. Bar plots indicate mean +/-se for following offspring groups: solid red Ddc +’/+, maternal WT; striped red Ddc -/+, maternal heterozygote; solid blue Ddc +/+’, paternal WT; striped blue Ddc +/-, paternal heterozygote. \**p* σ; 0.05.

**Figure S2.**
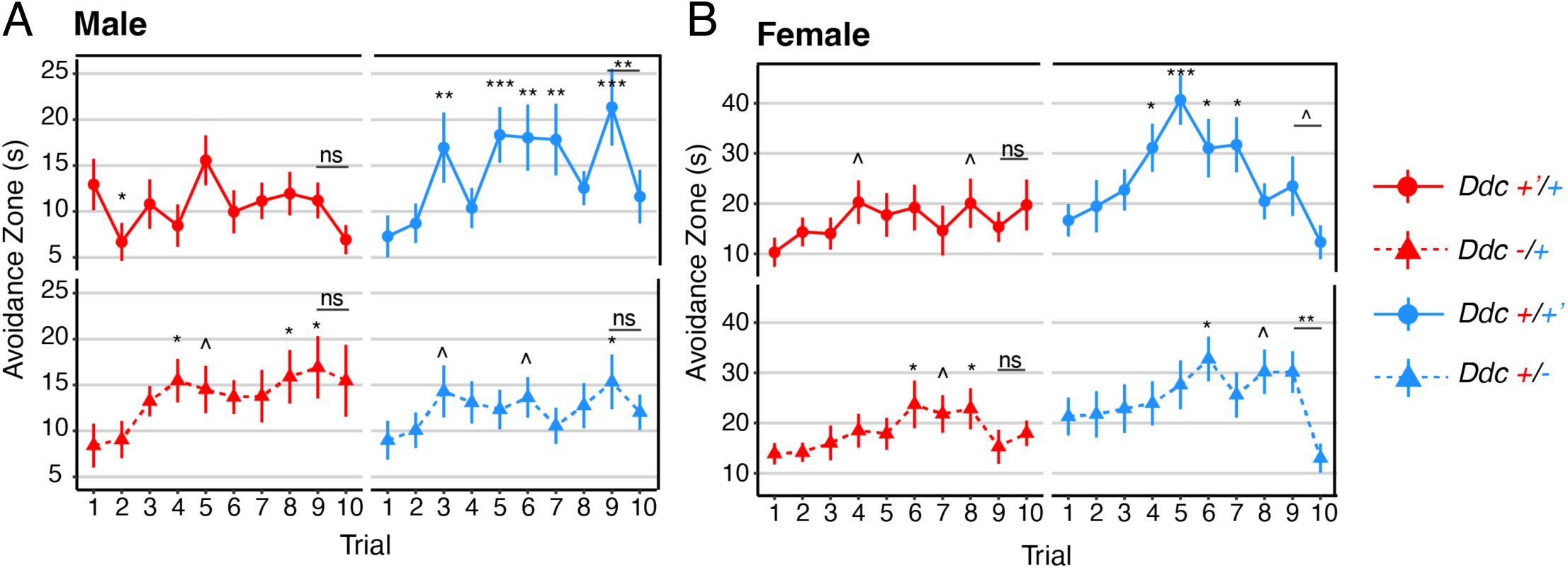
Avoidance zone time in social recognition test by trial and genotype for *Ddc*. **(a)** only *Ddc+/+’* paternal WT males showed a significant difference in time in avoidance zone (measured in seconds (s)) between trials 9 and 10. **(b)** only *Ddc+/-* paternal-null allele heterozygote females showed a significant differences in avoidance between trials 9 and 10. Point-range plots indicated mean +/-se values for each of the following groups: solid red line and circle, *Ddc* +’/+ maternal WT; dashed red line and triangle, *Ddc* -/+ maternal-null heterozygote; solid blue line and circle, *Ddc* +/+’ paternal WT; dashed blue line and triangle, *Ddc* +/-paternal-null heterozygote. Significant differences in Trials 2 – 9 were based on a comparison to Trial 1, significant differences in Trial 10 were based on a comparison to Trial 9 (indicated with a line below symbol): ^*p* ≤ 0.1, \**p* ≤ 0.05, \*\**p* ≤ 0.01, \*\*\**p* ≤ 0.001.

**Figure S3.**
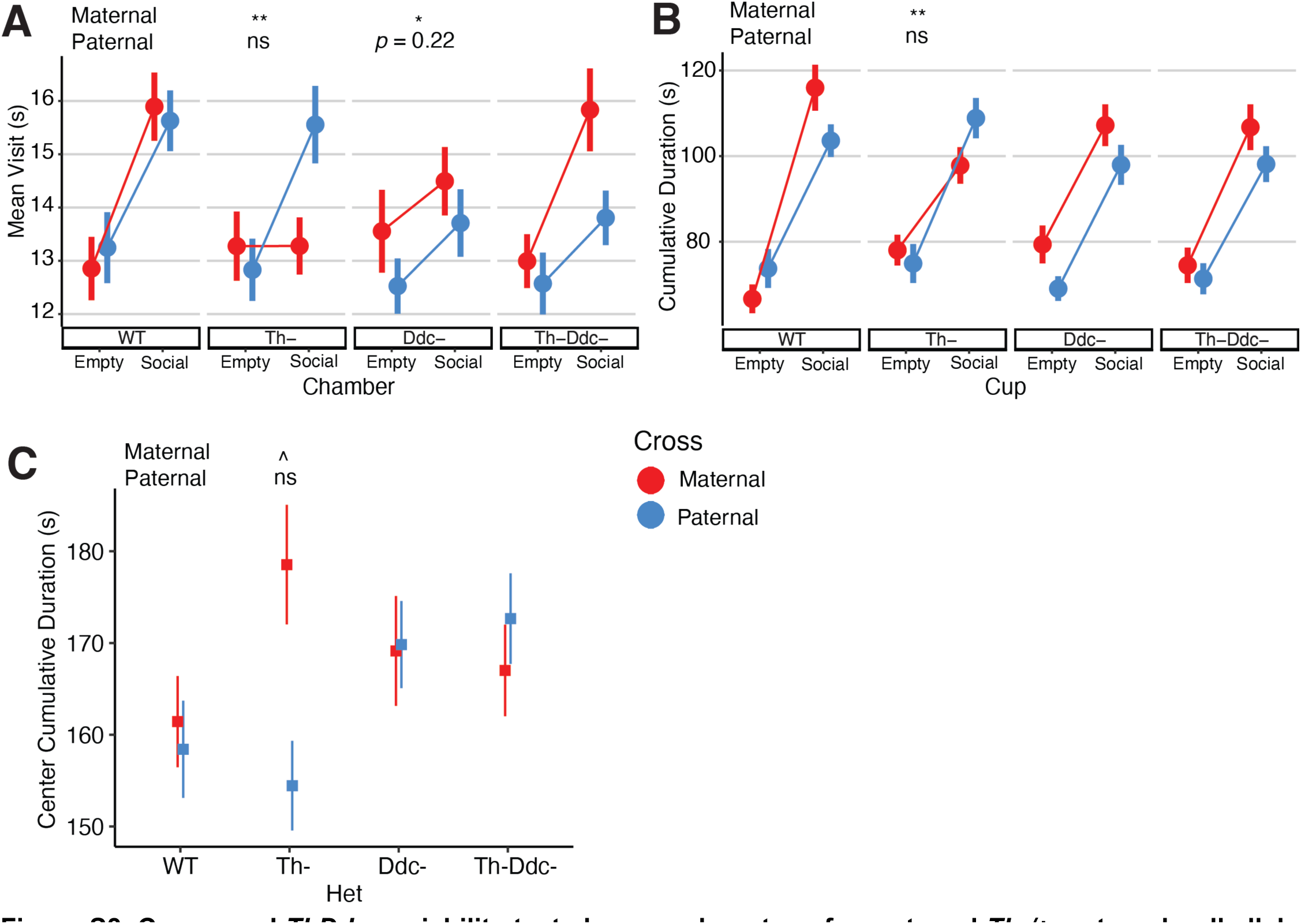
Compound *ThDdc* sociability test shows a phenotype for maternal *Th-/+* maternal-null allele heterozygote males. **(a)** Mean visit in seconds (s) duration for empty chamber and social chamber shows *Th-/+* maternal-null allele heterozygotes spend significantly shorter time visiting the social chamber than WT counterparts. **(b)** Cumulative duration of direct investigation of the cups shows maternal *Th-/+* maternal-null allele heterozygotes spend less time that WT counterparts sniffing the social cup. **(c)** Cumulative duration in center chamber also reveals a phenotype for maternal *Th-/+* heterozygotes. Point-range plots indicated mean +/-se values for each of the following groups: red, maternal cross offspring; blue, paternal cross offspring; Th-, *Th* hets.; Ddc-, *Ddc* hets.; Th-Ddc-, compound *Th* and *Ddc* hets. Significance symbols above heterozygous group data indicate multiple comparison adjusted post-test differences from maternal (top) and paternal (bottom) WT littermates: not significant (ns), ^*p* ≤ 0.1, \**p* ≤ 0.05, \*\**p* ≤ 0.01.

**Figure S4.**
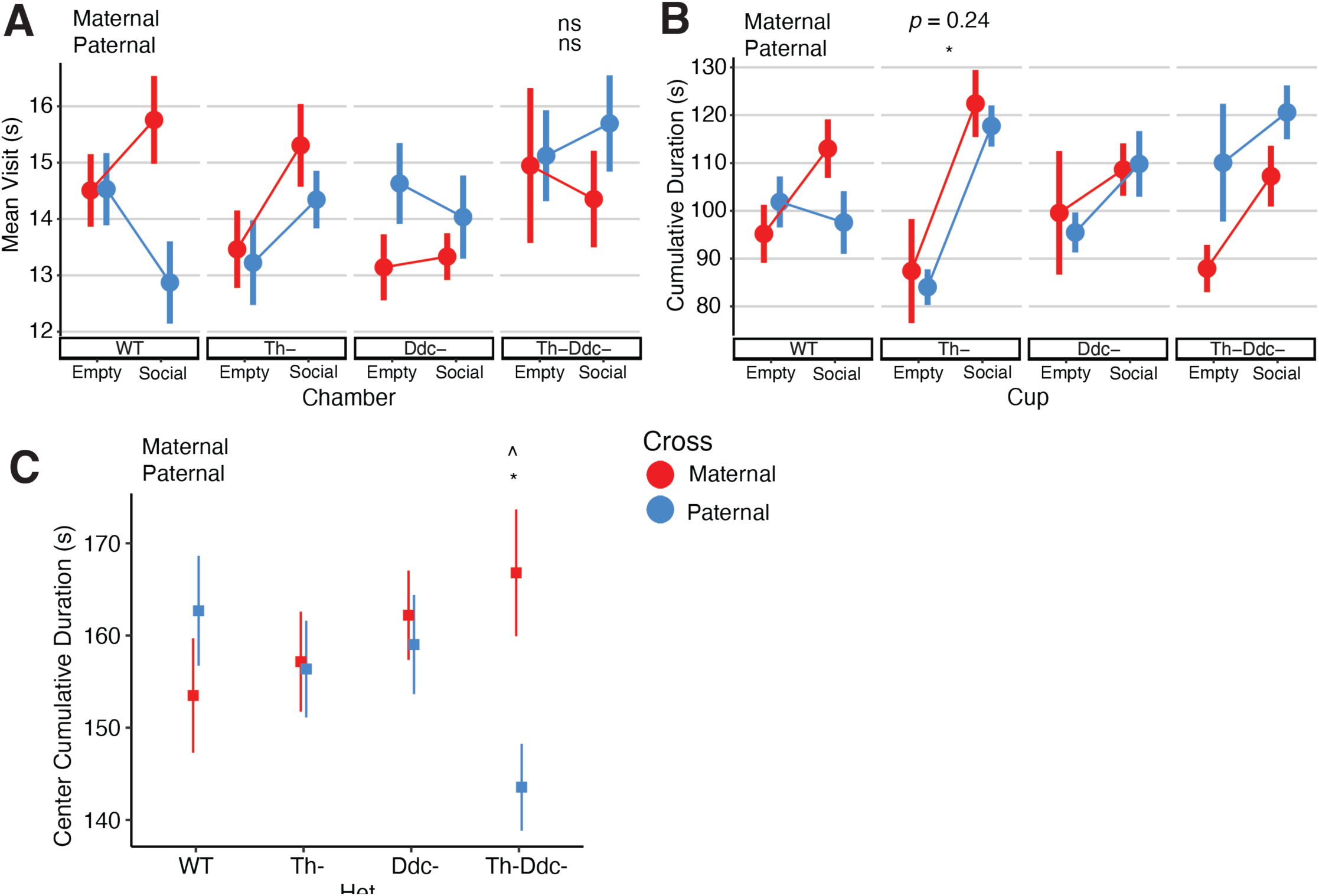
Additional measures of sociability in compound *ThDdc* females. **(a)** Mean visit duration in seconds (s) for empty chamber and social chamber. **(b)** Cumulative duration of direct investigation of the cups. **(c)** Cumulative duration in center chamber. Point-range plots indicated mean +/-se values for each of the following groups: red, maternal cross offspring; blue, paternal cross offspring; Th-, *Th* hets.; Ddc-, *Ddc* hets.; Th-Ddc-, compound *Th* and *Ddc* hets. Significance symbols above heterozygous group data indicate multiple comparison adjusted post-test differences from maternal (top) and paternal (bottom) WT littermates: not significant (ns), ^*p* ≤ 0.1, \**p* ≤ 0.05.

**Figure S5.**
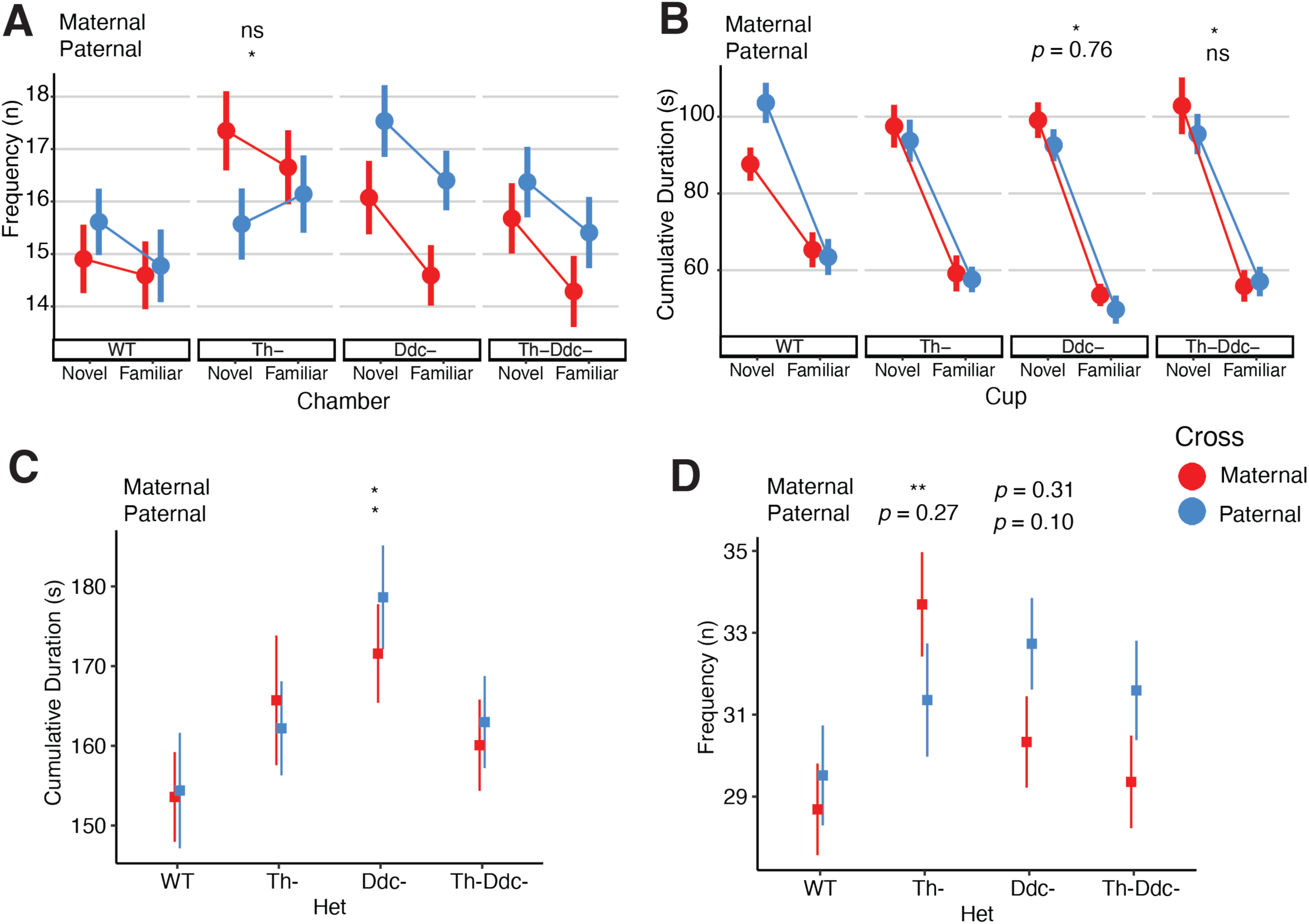
Additional measures of social novelty in compound *ThDdc* males. **(a)** Frequency (n) of chamber visits. **(b)** Cumulative duration of direct investigation of the cups in seconds (s). **(c)** Cumulative duration in center chamber. **(d)** Frequency of visits to center chamber. Point-range plots indicated mean +/-se values for each of the following groups: red, maternal cross offspring; blue, paternal cross offspring; Th-, *Th* hets.; Ddc-, *Ddc* hets.; Th-Ddc-, compound *Th* and *Ddc* hets. Significance symbols above heterozygous group data indicate multiple comparison adjusted post-test differences from maternal (top) and paternal (bottom) WT littermates: not significant (ns), ^*p* ≤ 0.1, \**p* ≤ 0.05, \*\**p* ≤ 0.01.

**Figure S6.**
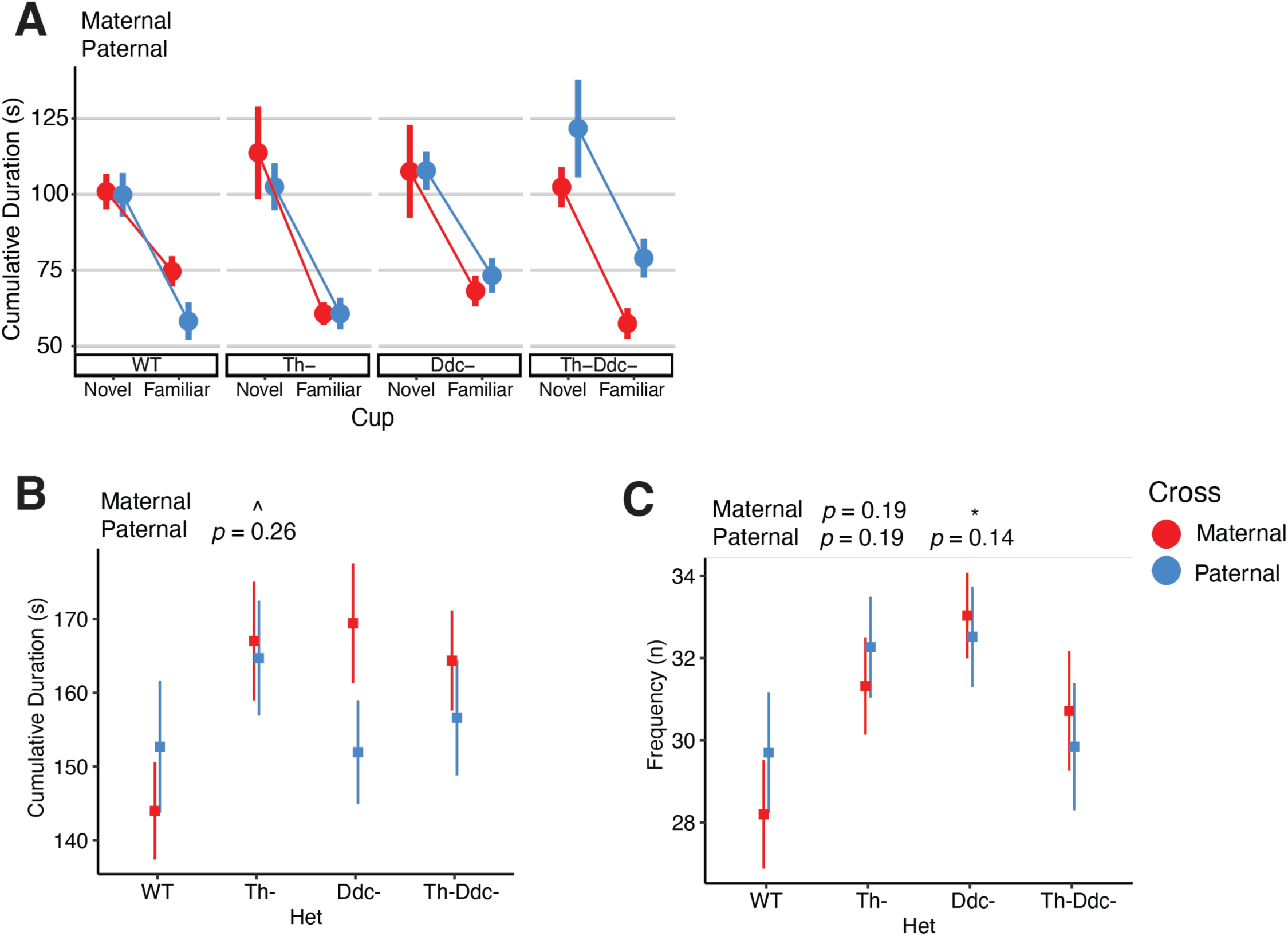
Additional measures of social novelty in compound *ThDdc* females. **(a)** Cumulative duration of direct investigation of the cups in seconds (s). **(b)** Cumulative duration in center chamber. **(c)** Frequency (n) of visits to center chamber. Point-range plots indicated mean +/-se values for each of the following groups: red, maternal cross offspring; blue, paternal cross offspring; Th-, *Th* hets.; Ddc-, *Ddc* hets.; Th-Ddc-, compound *Th* and *Ddc* hets. Significance symbols above heterozygous group data indicate multiple comparison adjusted post-test differences from maternal (top) and paternal (bottom) WT littermates: not significant (ns), ^*p* ≤ 0.1, \**p* ≤ 0.05, \*\**p* ≤ 0.01.

